# Non-uniform filament turnover and mechanically-driven contractility and bundle formation in disordered actomyosin networks

**DOI:** 10.1101/2025.11.05.686892

**Authors:** Alexander K. Y. Tam, Alex Mogilner, Dietmar B. Oelz

## Abstract

Bundles of actin filaments with similar positions and orientations commonly occur in cytoskeletal networks. We use mathematical modelling and simulation to investigate how filament turnover and mechanics influence contractility and bundle formation in disordered actomyosin networks. Using a two-dimensional agent-based model for an actomyosin network, we investigate four simplified models for filament turnover: uniform, biased, branching, and treadmilling. With no turnover, over time contractility decreases and bundle formation increases, and networks eventually form stationary patterns that cannot contract. Introducing turnover allows contractility to persist longer compared to the no-turnover scenario. Uniform turnover, where new filaments have random positions and orientations, disrupts bundle formation and enables persistent contractility. In biased turnover, branching, and treadmilling, the positions and orientations of new filaments depend on the existing network. These non-uniform turnover models increase bundle formation compared to uniform turnover, while still allowing long-term contractility. Branching at 70° disrupts bundle formation to enable prolonged contractility, whereas filament treadmilling disrupts the trade-off between bundle formation and contractility. Biased turnover places new filaments near existing ones, which promotes bundle formation but is less effective at maintaining contractility. We also varied mechanical factors in our simulations, especially filament bending flexibility and protein friction, which enhance bundle formation and contractility, respectively. Our results suggest that variations in turnover and mechanics might allow cells to tune contractility and bundle formation in disordered actomyosin networks.

## 1 Introduction

The actin cytoskeleton is a network of actin filaments (F-actin) and actin-binding proteins (ABPs) beneath the membrane and in the bulk of of biological cells. Interactions between F-actin and myosin-II minifilaments form actomyosin networks, which generate the mechanical forces underpinning cell shape, movement, and division [1–3]. Actin monomers (G-actin) form filaments (F-actin) with lengths on the micron scale [4–6]. Actin filaments are polar, with distinct plus (or barbed) and minus (or pointed) ends. Myosin-II minifilaments are molecular motors that bind to F-actin filament pairs and generate force and movement towards filament plus-ends [7]. Actin filaments are often also connected by cross-linker proteins that restrict relative motion between filaments, but do not move actively along them [8]. This creates a dense, cross-linked F-actin network, on which myosin-motor movement remodels the network and gives rise to contractile forces [9–12]. Throughout this work, we will refer to actin filaments (F-actin) as filaments, and myosin-II mini-filaments as motors.

Actomyosin networks were first studied in the context of muscle contraction. In muscles, filaments form linear arrays called sarcomeres, which have filament minus-ends in the centre and plus-ends pointed outwards and cantilevered into rigid Z-disks [13]. Relative motion of motors towards the filament plus-ends then pulls filaments inwards, generating contraction. This sliding filament mechanism was discovered decades ago [7], and the mechanism for how filament and motor mechanics generate muscle contraction is well-understood [13]. However, sarcomeric structure is not a necessary condition for actomyosin contraction. Sliding filament theory does not fully explain contraction in non-muscle cells, in which filaments are oriented and distributed seemingly at random. Many actomyosin networks contain stress fibres, which are extended cables of similarly-aligned filaments connected by cross-linker proteins [14–17]. Since filaments in stress fibres are aligned with alternating plus-ends and minus-ends [16, 18], they contract by a similar sliding filament mechanism to muscle cells [15]. Cells containing stress fibres produce stronger contraction than those without [14], highlighting that filament bundles are generally important for non-muscle cell contraction [19].

Mathematical modelling and simulations have been applied in previous studies of the actin cytoskeleton. Agent-based models for the movement of and interactions between individual proteins are common. Example agent-based models include the publicly-available software Cytosim [20], AFINES [21], and MEDYAN [22]. Modelling and simulation have been used to investigate the determinants of actomyosin contraction [12, 23–39]. These studies have shown that actomyosin contractility depends on factors including filament geometry [27, 32, 34], filament bending [28, 40], buckling [35, 37], protein friction [28, 40], cross-linking [26, 31], delayed myosin unbinding [23], drag forces [28], and filament treadmilling [12]. Other studies have focused on bundle formation in actomyosin networks [41–44]. Cross-linking [45–48] and attractive forces [49, 50] between filaments are key mediators of bundle formation in networks without myosin. Recent computational studies of bundle formation in actomyosin networks have demonstrated how cross-linking [22, 43, 51–53], initial orientation [52, 54, 55], motor density [52, 54, 56] and kinetics [57], and treadmilling [52, 54] affect bundle formation.

Understanding the mechanisms of actomyosin bundle formation in disordered networks is an open problem. Persistent contractile bundles do not readily emerge from standard models of disordered actomyosin networks. Many previous simulation studies have shown that disordered cytoskeletal networks eventually form stationary patterns, for example asters [11, 28, 58, 59]. These aster-like patterns have been observed during *in vitro* experiments with purified F-actin, non-muscle myosin-II, and cross-linker proteins [56, 58, 60]. Filament turnover disrupts these stationary patterns [61], and is a key feature in simulations that produce prolonged contractility [6, 40, 53, 62]. Turnover refers to the continuous exchange of network components between the network and the surrounding cytoplasm [3, 9, 63–65]. In cells, specialised mechanisms initiate filament turnover [65]. Although these turnover mechanisms are conserved across different cell types and organisms, their molecular complexity enables differential tuning in turnover processes (*e*.*g*. remodelling, severing, depolymerisation) and their rates [65]. We condense the complex molecular mechanisms into four simplified turnover models: uniform, biased, branching, and treadmilling. In uniform turnover, newly-recruited filaments have random positions and orientations. Unlike uniform turnover, in biased, branching, and treadmilling turnover the positions and orientations of new filaments depend on the existing network architecture.

In this manuscript, we investigate how mechanics and filament turnover affect contractility and bundle formation in disordered actomyosin networks. We adapt the mechanical model for filaments and motors of Tam, Mogilner, and Oelz [28], which is a system of force-balance equations incorporating filament mechanics (bending, drag, inextensibility, and protein-friction forces), motor inextensibility, and active motor movement. We simulate network evolution with four different models of filament turnover: uniform, biased, branching, or treadmilling. Our variational numerical scheme enables us to quantify contractility, and we introduce statistical indices to quantify filament aggregation and bundle formation. We then vary turnover rates, turnover models, and mechanical parameters in the simulations, to investigate their effects on contractility and bundle formation. Without turnover, filaments form stationary bundles that lose contractility. In general, non-uniform turnover increases bundle formation compared to uniform turnover, while still enabling persistent contractility. Branching prolongs contractility by disrupting bundle formation, whereas treadmilling lessens the trade-off between bundle formation and contractility. Mechanical factors, especially filament flexibility and protein friction, also promote contractility and bundle formation.

## 2 Materials and methods

We use a two-dimensional agent-based model to investigate how turnover and mechanics influence actomyosin bundle formation. We simulate the model using code written in Julia, quantifying both network contractility and bundle formation.

### 2.1 Mechanical model

Our agent-based model represents filaments and motors as individual entities [28, 34]. We represent filaments as chains of nodes connected by stiff linear springs, each spring having the same equilibrium length. Filaments are semi-flexible [66], and experience small-but-significant bending deformation. We represent motors as two nodes (motor domains) connected by a spring with equilibrium length zero. Motor nodes attach to filament pairs at their intersection, and each node moves actively towards the plus-end of the attached filament according to a linear force–velocity relationship [67]. The degrees of freedom of our model are the vector-valued filament-node positions, and the relative positions of motors along the filaments.

We do not explicitly model the movement of unattached motors or motors attached to a single filament. Instead, we remove motors from the network at a rate given by Bell’s law [68] to simulate force-dependent unbinding, and also remove a motor if either motor domain reaches the plus-end of a filament. When a motor is removed, we immediately replace the removed motor with a new motor located at an intersection between a pair of filaments. We select the filament–filament intersection at which to place the new motor randomly. While this approximation is relatively restrictive, our approach to motor binding and unbinding maintains the total number of motors throughout the simulation, preventing the number of motors from influencing the results.

Our model includes protein friction at all intersections between pairs of filaments that do not have an attached motor. We model protein friction as point-wise viscous drag that penalises relative motion between two intersecting filaments at their intersection. In practice, protein friction might represent cross-linker protein attachment to filament pairs [10], or solid friction between two filaments in contact [69]. Protein-frictional forces are larger than hydrodynamic-frictional forces between filaments and the cytoplasm, and have similar magnitude to forces exerted by motors [69]. Schematics of the filaments, motors, their interactions, and the relevant mechanical forces are illustrated in Figure 1.

**Figure 1:**
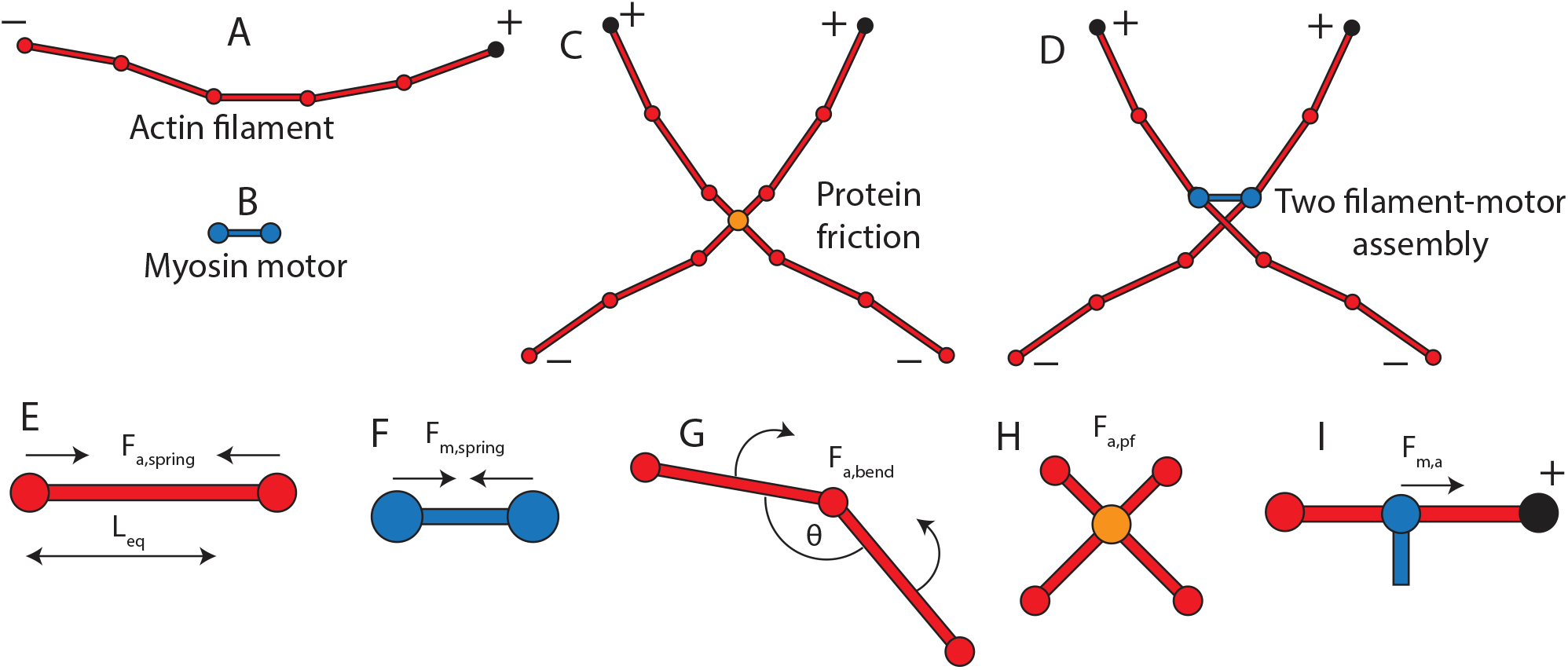
Schematic of filaments, motors, and their interactions (A–D), and mechanical features included in the model (E–I). (A) Diagram of filament as a series of nodes connected by linear springs (straight line segments) (B) Diagram of motor as two nodes connected by a spring. (C) Two-filament assembly incorporating protein friction. (D) Two-filament–motor assembly, with no protein friction. (E) Spring force, *F*_*a*,spring_, on a filament segment with equilibrium length *L*_eq_. (F) Spring force, *F*_*m*,spring_, on a motor (equilibrium length zero). (G) Bending force, *F*_*a*,bend_, on a pair of filament segments. (H) Protein friction, *F*_*a*,pf_, which restricts relative motion at filament intersections. (I) Active motor movement towards filament plus-ends, *F*_*m,a*_.

We express our mechanical model as the system of force-balance equations,

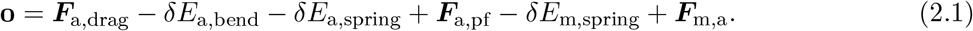

The first four terms on the right-hand side of (2.1) are the mechanical contributions of the filaments. The symbol ***F***_a,drag_ represents viscous-drag forces between the filaments and the cytoplasm. The filament bending force is the variation of the filament bending energy *E*_a,bend_, where *E*_a,bend_ integrates elastic potential energy along the extent of a filament. We model the filament stretching force as the variation of *E*_a,spring_, which is the sum of spring forces for each segment between adjacent nodes of a filament. We assume that each spring satisfies Hooke’s law, with spring constant *k*_a_. Since filaments are effectively inextensible [70], we assume a large value of bending stiffness, *k*_a_ = 1000 pN nm^*−*1^, for every filament segment. The symbol ***F***_a,pf_ represents viscous-drag forces due to protein friction. These drag forces apply point-wise at intersections between filaments, and penalise relative motion of the intersecting filaments.

The final two terms on the right-hand side of (2.1) are mechanical contributions from motors. We represent motor-stretching forces as the variation of spring energy *E*_m,spring_. Like filaments, we assume that a spring satisfying Hooke’s law connects the two motor nodes. We assume this spring to be stiff, with spring constant *k*_m_ = 1000 pN nm^*−*1^, to maintain a motor length close to zero throughout the simulation. The final force term, ***F***_m,a_, models filament–motor interactions, for which we adopt a linear force–velocity relationship. Like the model of Tam, Mogilner, and Oelz [28], we neglect thermal forces, which can hasten bundle formation but do not affect the eventual morphology [45].

### 2.2 Turnover models: uniform, biased, branching, and treadmilling

Turnover refers to the continuous exchange of network components between the network and cytoplasm [6]. To maintain the total number of filaments throughout a simulation, we implement filament turnover by removing a filament (and any motors attached to the filament) from the network, and immediately replacing the removed filament (and motor, if applicable) with a new one. Removal of filaments from the simulated network might represent filament disassembly by severing [71, 72], and filament addition might represent filament nucleation [71]. In this work, we develop methods to mimic filament turnover without explicitly modelling severing, the positions of actin monomers and nucleation complexes, or polymerisation and depolymerisation of the filament ends. Simulation models often consider turnover whereby the position and orientation of new filaments is random [6, 28, 40]. However, severing and nucleation rates depend on the density of ABPs such as ADF/cofilin and gelsolin [71, 73, 74], and the density of actin monomers. If ABPs are not uniformly distributed in space, filament turnover might exhibit spatial bias. Indeed, Goode, Eskin, and Shekhar [65] write that turnover involves a polarised flux of actin towards specific regions in the network, indicating spatial non-uniformity.

We implement four turnover models: uniform turnover, biased turnover, branching turnover, and treadmilling turnover. Regardless of turnover model, we assume that filaments undergo turnover with rate *k*_off,*a*_, assumed constant and the same for every filament. The probability that a filament turns over in one time step of the simulation is then 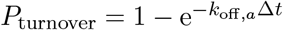, where Δ*t* is the time-step size. In uniform turnover, new filaments have zero curvature, and have positions and orientations chosen at random from uniform distributions (see Figure 2A). In contrast, the existing network architecture influences the positions and orientations of new filaments in biased, branching, and treadmilling turnover.

**Figure 2:**
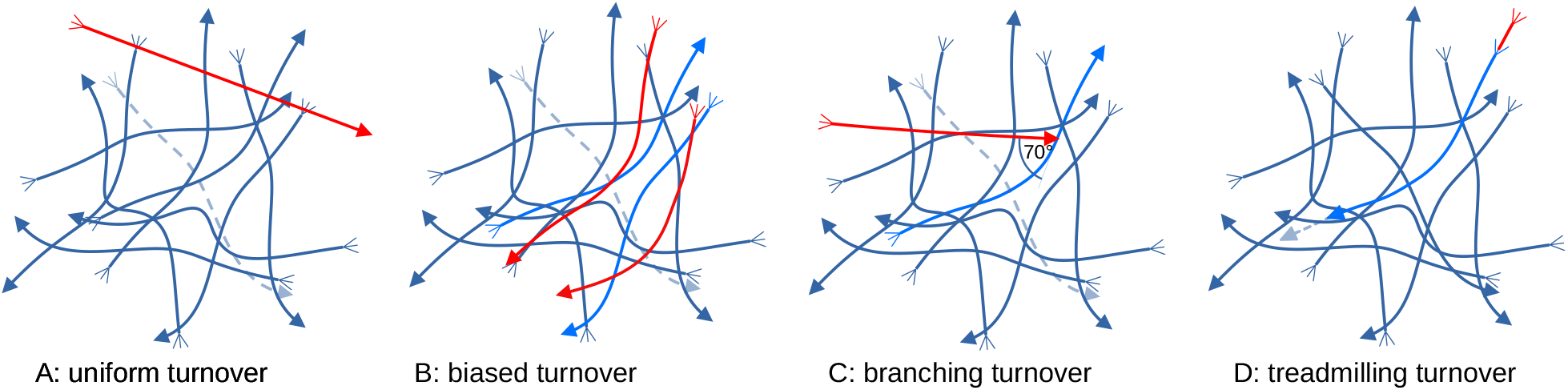
Illustration of the different models for filament turnover considered in this study. In each sketch, dark blue curves represent filaments (arrows indicate minus-ends and tridents indicate plus-ends). Dashed curves illustrate the random selection and removal of a filament from the network, and red curves represent newly-added filaments due to turnover. Where appropriate, light blue filaments represent a reference filament on which the newly-added filament is based. (A) Uniform turnover, where the new filament has random position and orientation and zero curvature. (B) Biased turnover, where the new filament is a copy of an existing filament, with a translation and rotation, and sometimes polarity reversal. (C) Branching turnover, in which new filaments appear at an angle of 70° to existing filaments. (D) Treadmilling turnover, where a filament depolymerises (shrinks) at its minus-end and polymerises (grows) at its plus-end.

Biased turnover captures a tendency for new filaments to form close to existing ones. In biased turnover, after removing a filament we introduce the new one by first copying an existing reference filament selected at random from those remaining in the network. We then rotate the newly-copied filament *t,θ* about its centre, by a normally-distributed random angle 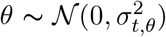, and translate the rotated filament by a vector 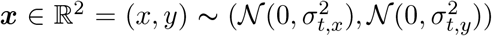, where ***x*** is dimensionless and scaled by domain width. The standard deviations *σ*_*t,θ*_, *σ*_*t,x*_, and *σ*_*t,y*_ are user-prescribed constants. Smaller standard deviations result in a stronger effect of the existing network architecture on turnover. When implementing biased turnover we also swap the plus-end and minus-end of the new filament with probability *P*_rev_. In typical simulations, we set *P*_rev_ = 0.5. Reversing the filament polarity mimics the observation that plus-ends and minus-ends alternate in stress fibres [18].

Filament branching mediated by the Arp2/3 complex [75] is another turnover scenario that we implement. By capping the filament minus-ends [75], Arp2/3 stimulates nucleation of new daughter filaments that grow at an angle of 70 *±* 7° from the side of the mother [75]. To implement branching, when we remove a filament from the network we introduce a new filament with zero curvature whose minus-end coincides with a random position along a randomly-selected reference filament. The new filament is oriented at a *θ*_*b*_ degree angle from the relevant segment of the reference filament. We simulate networks with different values of *θ*_*b*_, but *θ*_*b*_ = 70° corresponds to Arp2/3-mediated branching. Branching turnover is illustrated in Figure 2C.

Treadmilling represents actin remodelling by simultaneous plus-end polymerisation and minus-end depolymerisation. Polymerisation occurs when G-actin monomers associate with the end of an actin filament, and occurs more readily at the plus-end than at the minus end [76, 77]. Minus-end depolymerisation might occur due to cofilin, an ABP that catalyses remove actin monomers from the minus-end [17, 71]. To implement treadmilling, we remove the filament segment closest to the minus-end, and add a new segment to the filament at the plus-end (Figure 2D). We assume that new segments have the equilibrium segment length, and the same orientation as the adjacent segment.

### 2.3 Numerical methods

We implement the model in a spatially two-dimensional simulation environment written in Julia. Our code simulates network evolution using discrete, equispaced time steps. In each time step, we first simulate filament turnover and motor binding and unbinding. We then solve the force-balance equations (2.1) to implement the mechanical component of the model. We solve the mechanical model by obtaining the filament and motor positions that minimise the functional

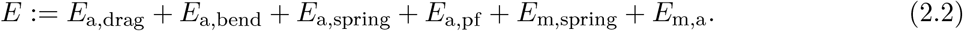

Each term in (2.2) corresponds to a term in the force-balance equation (2.1). The symbols *E*_a,drag_, *E*_a,pf_, and *E*_m,a_ denote terms whose variations are finite-difference approximations of the forces ***F***_a,drag_, ***F***_a,pf_, and ***F***_m,a_, respectively, which we interpret as dissipative energies in the context of Onsager’s variational principle [78]. In Julia, we minimise the energy functional (2.2) using the limited-memory Broyden–Fletcher–Goldfarb–Shanno (LBFGS) method [79–82], using the `Optim.jl` package [83].

### 2.4 Quantifying contractility

In our simulations, we assume that filament motion gives rise to uniform elongation and shearing of the two-dimensional domain in which the network resides. This assumption neglects the details of where forces are propagated through the network and their direction, and is common in actomyosin network simulations [5, 6, 12, 28]. Uniform deformation would occur for a dense network with numerous overlapping copies of the reference network of simulated filaments, as shown in Figure 3A. Such a dense network is a reasonable approximation of a real network. Under the assumption of uniform deformation in the domain, we can associate the forces occurring within the domain with uniform forces at the boundaries. We can then quantify contractility by viewing the domain as a simple plane stress element. We discuss this scenario further in the context of a two-filament–motor assembly in Tam, Mogilner, and Oelz [34].

**Figure 3:**
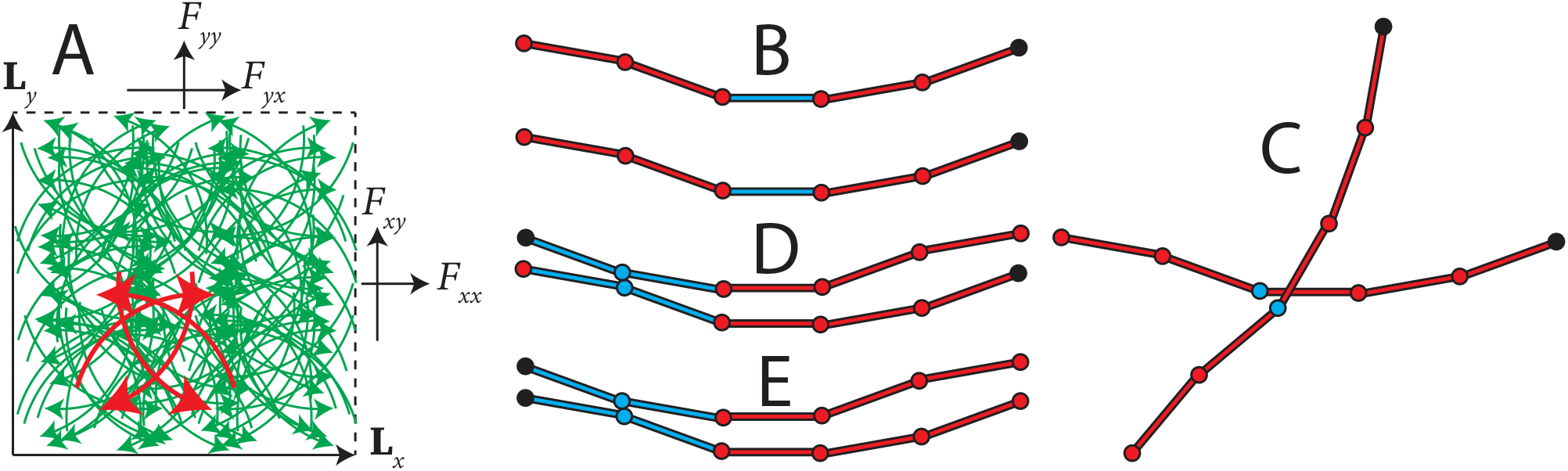
(A) Diagram of the simulation domain, which is the parallelogram spanned by the two vectors ***L***_*x*_ and ***L***_*y*_. The force vectors ***F***_*x*_ = (*F*_*xx*_, *F*_*xy*_) and ***F***_*y*_ = (*F*_*yx*_, *F*_*yy*_) contain the normal and shear forces acting on the domain boundaries. We assume the dense network consists of numerous overlapping copies of the reference (simulated) network highlighted in red, such that deformations are uniform throughout the domain. (B–E) Illustration of the statistical indices for aggregation, bundle formation, and parallel bundles (2.8). Red circles are nodes in our representation of filaments, and the black circle is the plus end. (B) Parallel filament pair with similar orientation, but not sufficiently close. This pair is not included in *N*_*a*_, *N*_*b*_, or *N*_*p*_. (C) Pair with nodes that are sufficiently close, but with dissimilar orientations. This pair is included in *N*_*a*_, but not *N*_*b*_ or *N*_*p*_. (D) Anti-parallel filament pair with nodes that are sufficiently close, and have similar orientation. This pair is included in *N*_*a*_ and *N*_*b*_, but not *N*_*p*_. (E) Parallel filament pair with nodes that are sufficiently close, and have similar orientation. This pair is included in *N*_*a*_, *N*_*b*_, and *N*_*p*_.

For uniform elongation and shear, we can use the energy minimisation technique to compute the forces on the domain boundary. We define

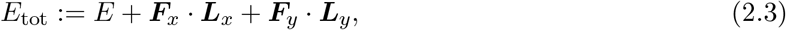

where *E* is defined in (2.2), and ***L***_***x***_ = (*L*_*xx*_, *L*_*xy*_) and ***L***_*y*_ = (*L*_*yx*_, *L*_*yy*_) are vectors specifying two edges of the domain. The vectors ***F***_*x*_ and ***F***_*y*_ contain the normal and shear force components on the domain boundary. These are illustrated in Figure 3A. In numerical simulations with constant domain size and shape, unconstrained minimisation of (2.3) is equivalent to minimising (2.2), where the force vectors ***F***_*x*_ and ***F***_*y*_ are Lagrange multipliers that enforce the constant-domain-size constraints. In numerical simulations, at each time step we compute force components using

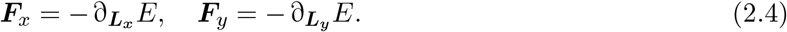

Neglecting possible out-of-plane stresses, the state of stress in the network is given by the two-dimensional stress tensor

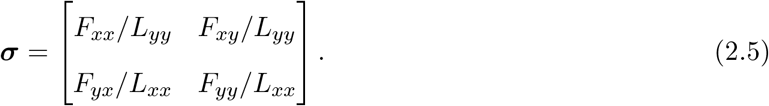

The mean normal stress,

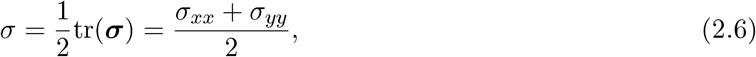

then provides a measure of instantaneous network contractility, and by convention negative *σ* indicates contraction, and positive *σ* indicates expansion. We also introduce the time-averaged mean normal stress,

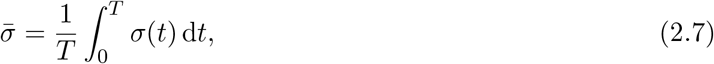

where *T* is the total simulation duration. The time-averaged mean normal stress 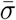 is our primary measure to quantify the net expansion or contraction generated in simulated networks. Throughout this work, we denote all quantities averaged over time using bars.

### 2.5 Quantifying aggregation and bundle formation

We define three statistical indices to quantify aggregation and bundle formation. These indices,

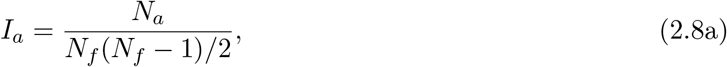

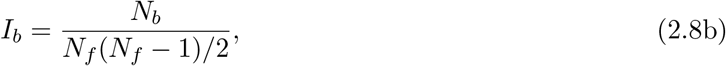

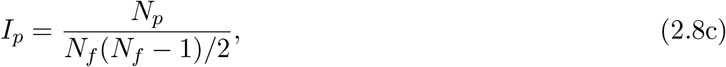

are the aggregation index, bundle index, and parallel-bundle index respectively. All three indices are ratios of the number of a particular type of filament pair, to the total number of pairs in the network, *N*_*f*_ (*N*_*f*_ *−* 1)*/*2, where *N*_*f*_ is the total number of filaments. To compute the aggregation index (2.8a), we count the number of filament pairs that are sufficiently close to each other, *N*_*a*_. We assume that two filaments are sufficiently close if any node on one filament is within 0.25 µm of any node of the other filament. In the bundle index (2.8b), *N*_*b*_ is the number of aligned close filament pairs in the network. A pair is included in *N*_*b*_ if the two filaments are sufficiently close, using the same definition as for *N*_*a*_, and have similar orientation. We classify a filament pair as having similar orientation if the acute angle between any pair of sufficiently close segments is less than or equal to 20°. Finally, the count *N*_*p*_ for the parallel-bundle index (2.8c) is similar to *N*_*b*_, but excludes antiparallel pairs where plus ends are located opposite to each other. Figure 3B–E illustrates the conditions for a filament pair to be included in *N*_*a*_, *N*_*b*_, and *N*_*p*_.

We compute the three indices (2.8) in simulations to measure the combinations of filament mechanics, turnover, and model parameters that give rise to bundle formation. Similar to the time-averaged mean normal stress (2.7), we also compute the time-averaged indices

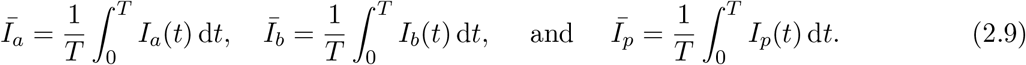

Our indices provide general statistics to quantify bundle formation, and apply to simulations with different turnover methods and turnover rates. For example, the index *I*_*b*_ provides a useful single measure of overall bundle formation within a network, but does not explicitly identify individual bundles. One example of the random initial condition used in our simulations gave the baseline values *I*_*a*_ = 0.1322, *I*_*b*_ = 0.0249, and *I*_*p*_ = 0.0130. Throughout this work, we can consider an index elevated if the value of the index exceeds the baseline value. Using these indices, we focus on network-scale contractility and bundle formation, rather than measuring the contractility within individual bundles.

## 3 Network simulation results

We obtain results for simulations with 150 filaments (each with equilibrium length of 1 µm) and 30 motors on a 5 µm *×* 5 µm domain, until *t* = 300 s using a time-step size of Δ*t* = 0.05 s. For each combination of parameters and turnover methods, we perform 10 simulations with a random initial condition. When reporting results, we denote quantities that are averaged across all simulations using angled bracket notation. For example, the trial-averaged mean normal stress is

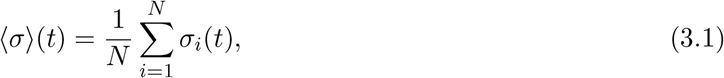

for each realisation *σ*_*i*_(*t*) for *i* = 1, … *N*, where *N* = 10. We denote time-averaged and trial-averaged quantities using both a bar and angled brackets. In subsequent sections, we often show results that are smoothed using a Savitzky–Golay filter, performed using the Loess Julia package with span parameter *α* = 0.2. Unless otherwise specified, parameters take the default values [4–6, 12, 25, 28, 56, 62, 64, 66, 69, 75, 84–91] indicated in the Supporting Material. Further explanation of these parameter values are provided in Tam, Mogilner, and Oelz [28]. Our key results are outlined in the subsections below.

### 3.1 Trade-off between bundle formation and contractility

Our model reproduces the known result that networks without turnover lose contractility over time [11, 28, 58, 59]. Figure 4 summarises simulation results with no turnover. Figure 4A displays the smoothed, time-averaged mean normal stress, showing that networks are initially contractile, but lose contractility after approximately *t* = 100 s to 150 s. Figure 4B shows that the aggregation, bundle formation, and parallel-bundle indices all increase initially, before reaching a plateau around the time that the network loses contractility. Increases in overall aggregation and bundle formation are largely explained by parallel-bundle formation, as myosin motors reorganise the network and draw filaments together. These parallel structures limit future contraction, because motors moving along two parallel filaments will not generate contraction [34]. These results with no turnover do not explain some bundles that remain highly contractile, such as stress fibres and the contractile ring. In those structures, filaments form sarcomeric arrangements with plus-ends linked together in periodic locations, and minus-ends overlapping in between [41, 42], or have plus-ends anchored to focal adhesions on the membrane [44]. Our disordered networks lack both of these features. Instead, the key mechanism for generating contractility in our simulations lateral filament bending [28], which is absent in the stationary bundles.

**Figure 4:**
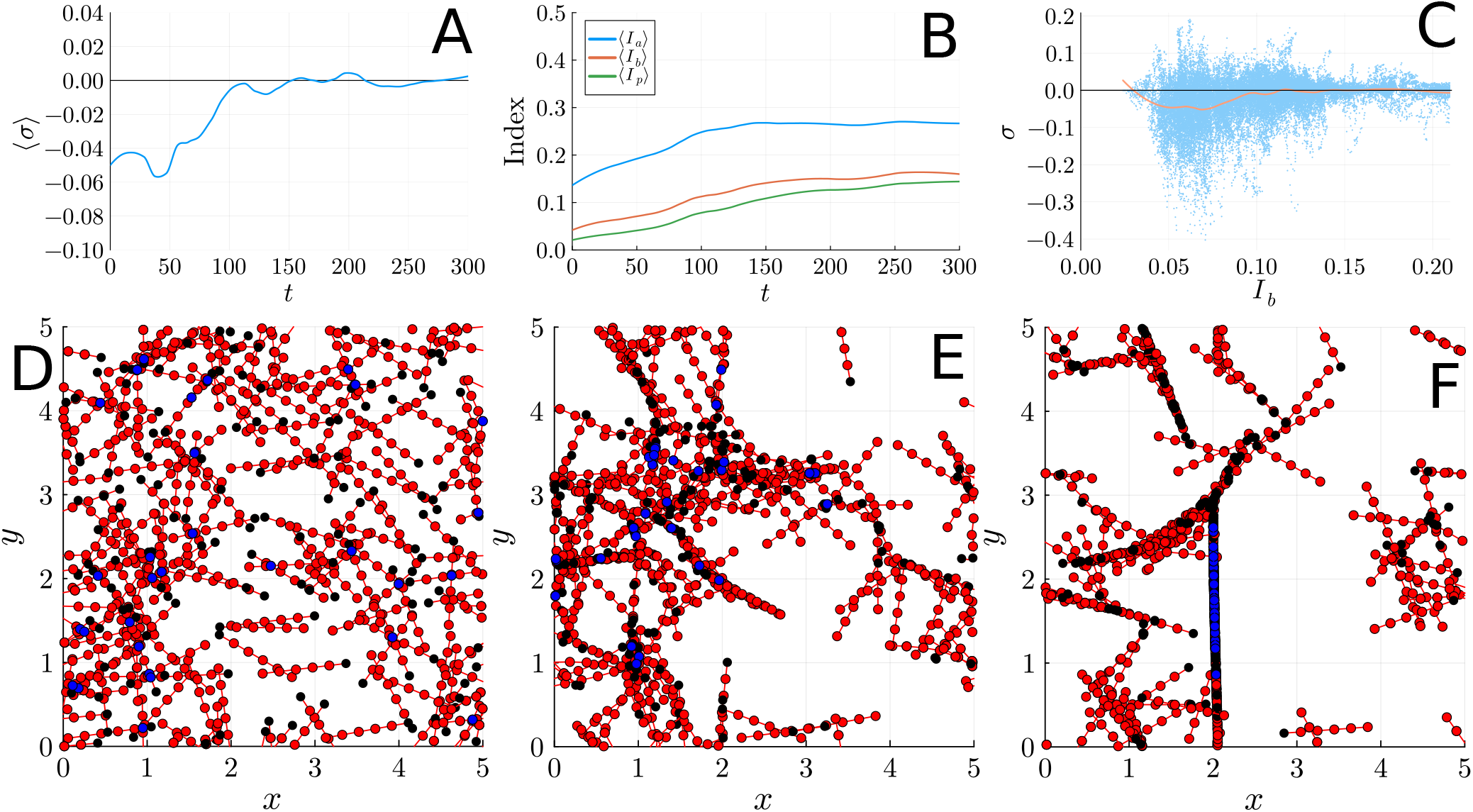
Numerical results for simulations with no turnover. (A) Trial-averaged mean normal stress over time for *N* = 10 simulations with no turnover, smoothed using a Savitzky–Golay filter (LOESS regression). (B) Trial-averaged bundle indices over time for *N* = 10 simulations with no turnover, smoothed using a Savitzky–Golay filter (LOESS regression). (C) Comparison of instantaneous measurements (blue dots) of mean normal stress and bundle index across all simulations and time. The solid curve is a LOESS regression of the data. (D) Example network configuration at *t* = 0. (E) Configuration of the same network at *t* = 50. (E) Configuration of the same network at *t* = 300.

Figure 4C shows the relationship between bundle index and stress, showing that networks with bundle indices *I*_*b*_ *<* 0.1 are contractile, but increased bundle formation leads to loss of contractility. The sample networks in Figure 4D–F reflect that excessive bundling inhibits contractility. The network is initially disordered (see Figure 4D), but after a period of initial contraction distinct parallel structures form, increasing bundle formation but preventing sustained contractility. However, contractility also decreases if *I*_*b*_ is low, which can occur if filaments are not aggregated, reducing filament–filament interactions. This result suggests that an intermediate amount of bundle formation, approximately *I*_*b*_ = 0.07 for the scale of the simulated networks, is optimal for contractility. The amount of bundling that is optimal for contractility is 2–3 the amount of bundling in the random initial condition, *I*_*b*_ = 0.0249.

### 3.2 Uniform turnover prevents bundling and supports long-term contractility

Uniform turnover influences the trade-off between contractility and bundle formation by disrupting bundles, enabling prolonged contractility. This effect of uniform turnover has also been reported in previous studies [6, 40, 53, 62]. Figure 5 summarises the results for uniform turnover with rate *k*_off,*a*_ = 0.04 s^*−*1^, a turnover rate estimated for cortical actomyosin networks [28]. This value of *k*_off,*a*_ enables comparison with the no-turnover results in Figure 4. Figure 5A shows that with uniform turnover the networks maintain contractility, with similar magnitude to the initial period of contraction of a disordered network. Figure 5B and Figure 5C suggest that the prolonged contractility occurs because uniform turnover disrupts bundle formation. In Figure 5B, there is little change in the aggregation, bundle formation, or parallel-bundle formation indices over time. Figure 5C shows a similar relationship between bundle formation and contractility to Figure 4C, but due to turnover the amount of bundle formation never becomes sufficiently high to prevent contractility. Instead, *I*_*b*_ remains sufficiently low, within the range of values for *I*_*b*_ that Figure 4C and Figure 5C indicate are contractile. Figure 5D–F also do not show parallel-bundle formation similar to Figure 4D–F as the network evolves, reinforcing the idea that uniform turnover disrupts parallel bundles.

**Figure 5:**
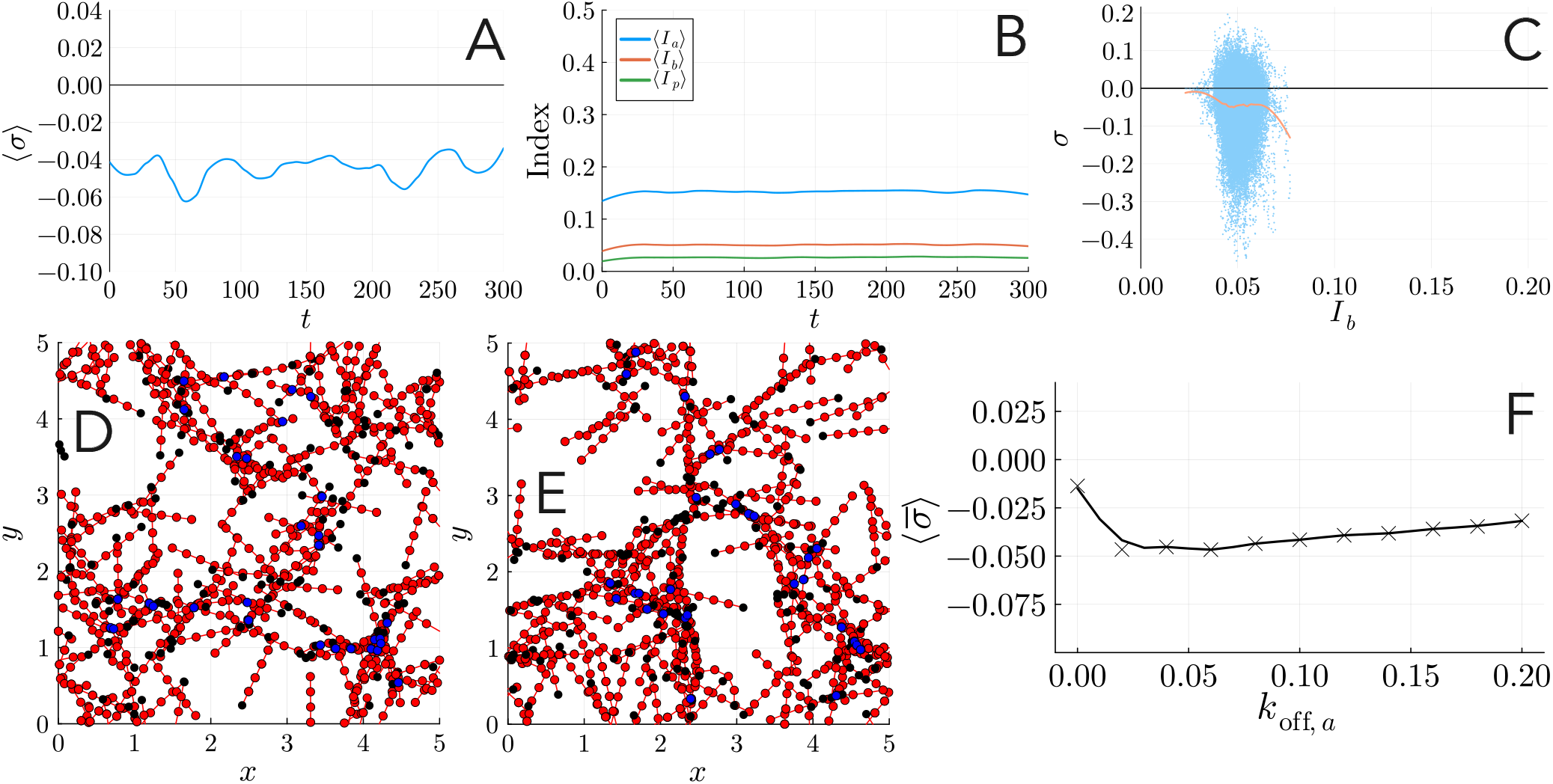
Numerical results for simulations with uniform turnover with rate *k*_off,*a*_ = 0.04 s^*−*1^. (A) Trial-averaged mean normal stress over time for *N* = 10 simulations, smoothed using a Savitzky– Golay filter (LOESS regression). (B) Trial-averaged bundle indices over time for *N* = 10 simulations, smoothed using a Savitzky–Golay filter (LOESS regression). (C) Comparison of instantaneous measurements (blue dots) of mean normal stress and bundle index across all simulations and time. The solid curve is a LOESS regression of the data. (D) Example network configuration at *t* = 50. (E) Configuration of the same network at *t* = 300. (F) Relationship between uniform turnover rate and contractility. The solid curve is a LOESS regression of the data.

In the fast-turnover rate limit, the network will approach the level of bundle formation associated with a random initial condition, *I*_*b*_ *≈* 0.0249. This value of *I*_*b*_ is less than the amount of bundle formation that maximises contractility, as shown in Figure 4C and reinforced in Figure 5C. Simulations with varying turnover rates suggest that there is an optimal uniform-turnover rate lying between *k*_off,*a*_ = 0.02 s^*−*1^ and *k*_off,*a*_ = 0.06 s^*−*1^ that maximises contractility. Interestingly, the turnover rate in cortical networks, *k*_off,*a*_ *≈* 0.04 s^*−*1^, is close to optimal for generating contraction in our simulations. The relationship between uniform-turnover rate and contractility is shown in Figure 5F, and more detailed results are in the Supporting Material.

### 3.3 Biased turnover increases bundle formation and leads quickly to loss of contractility

Since the positions and orientations of new filaments depend on the existing network in the non-uniform turnover models, with non-uniform turnover the amount of filament reshuffling will be more than with no turnover, but less than with uniform turnover. In biased turnover, new filaments are placed close to existing filaments. As Figure 6A shows, contractility decreases quickly before *t* = 50 s, before settling in a seemingly steady state of weak contractility. Unlike the no-turnover scenario, the presence of turnover means that the network does not form stationary patterns and lose contractility completely. However, the spatial bias means that the contractility with biased turnover is much less than the contractility with uniform turnover, for the same turnover rate.

**Figure 6:**
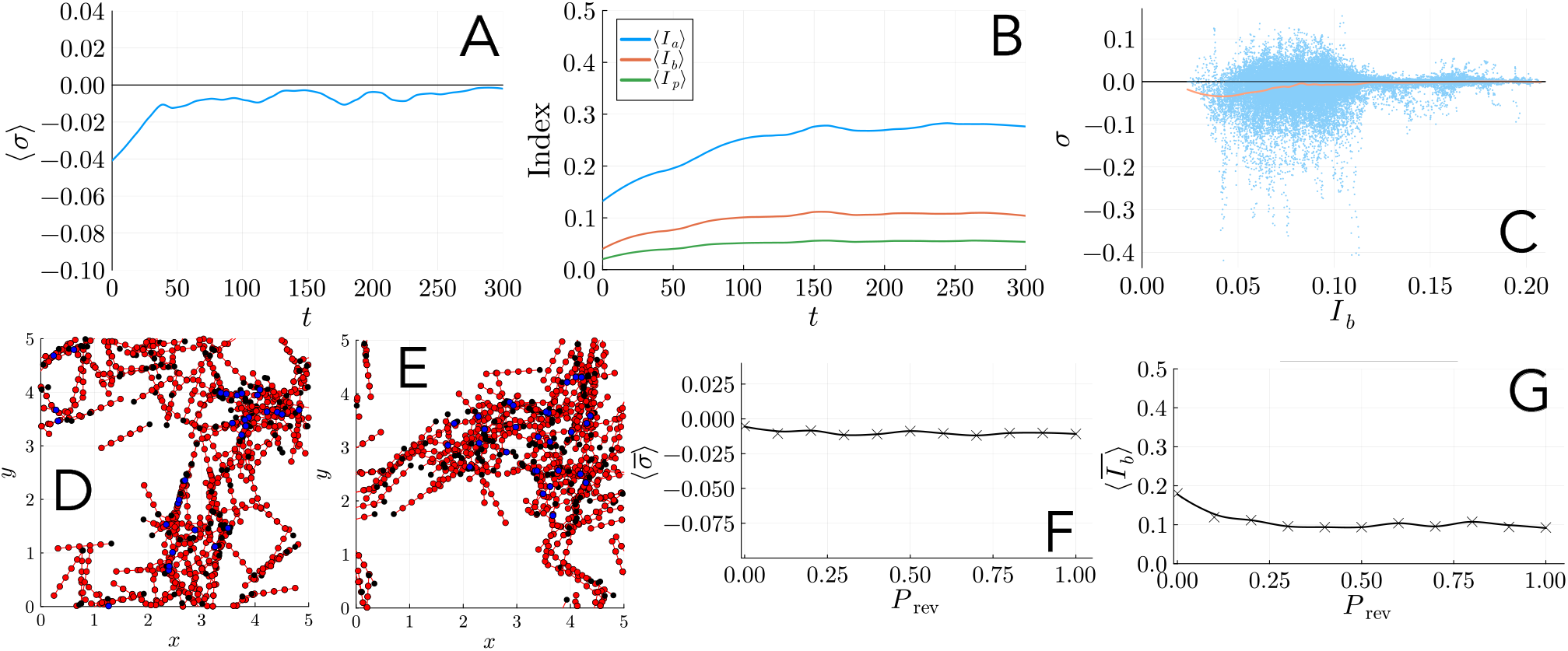
Numerical results for simulations with biased turnover with rate *k*_off,*a*_ = 0.04 s^*−*1^. (A) Trial-averaged mean normal stress over time for *N* = 10 simulations, smoothed using a Savitzky– Golay filter (LOESS regression). (B) Trial-averaged bundle indices over time for *N* = 10 simulations, smoothed using a Savitzky–Golay filter (LOESS regression). (C) Comparison of instantaneous measurements (blue dots) of mean normal stress and bundle index across all simulations and time. The solid curve is a LOESS regression of the data. (D) Example network configuration at *t* = 50. (E) Configuration of the same network at *t* = 300. (F) Impact of the polarity reversal probability, *P*_rev_, on contractility. (G) Impact of the polarity reversal probability, *P*_rev_, on bundle formation.

Figure 6B shows how biased turnover affects aggregation and bundle formation. All three indices *I*_*a*_, *I*_*b*_, and *I*_*p*_ increase over time. Compared to simulations with no turnover, biased turnover yields similar aggregation (*I*_*a*_), but less bundle formation (*I*_*b*_) and parallel-bundle formation (*I*_*p*_). The trends in the bundle index, *I*_*b*_, are consistent with the idea that contractility decreases as bundle formation increases. The relationship between *I*_*b*_ and *σ* shown in Figure 6C, which is similar to Figure 4C, reinforces this idea. With biased turnover, the bundle index stabilises at *I*_*b*_ *≈* 0.1, which is just low enough to maintain weak contractility.

With biased turnover, the difference *I*_*b*_ *−I*_*p*_ also increases over time, suggesting an increase in antiparallel bundles. This trend is due to the possibility of polarity reversal in biased turnover, and does not occur with no turnover or uniform turnover. Without turnover, the motors will instead pull filaments into stationary parallel arrangements. This difference is also reflected in the network patterns, as Figure 6E shows. The network with biased turnover forms a thicker aggregate, rather than a thin stationary pattern like those in Figure 4F. If polarity reversal is prevented, *σ* and *I*_*b*_ become more similar to the case of no turnover, with less contractility and more bundle formation. This is shown by the left-most points in Figure 6F and Figure 6F. Introducing even a small amount of polarity reversal, for example *P*_rev_ = 0.1, is sufficient to increase contractility and decrease overall bundle formation. Further details on the effects of *P*_rev_ are available in the Supporting Material.

### 3.4 70° branching favours contractility over bundle formation

Figure 7 shows how Arp2/3-mediated filament branching affects contractility and bundle formation. Like biased turnover, networks with branching lose some contractility within the first 50 s, before settling at an approximately steady level of contractility (see Figure 7A). However, networks with branching turnover are more contractile than those with biased turnover. A possible explanation for this increased contractility is that branching is more effective than biased turnover at disrupting bundle formation. As Figure 7B shows, the amount of aggregation with branching is similar to biased turnover. The network snapshot in Figure 7F illustrates this result, where a filament aggregate forms on the right-hand side of the domain. However, the networks with branching maintain less overall bundle formation compared to biased turnover (Figure 7B). This reduced bundling enables networks with branching to remain more contractile than networks with no turnover or biased turnover. However, branching remains less effective at promoting contractility than uniform turnover, because the positions and orientations of new filaments exhibit spatial bias.

**Figure 7:**
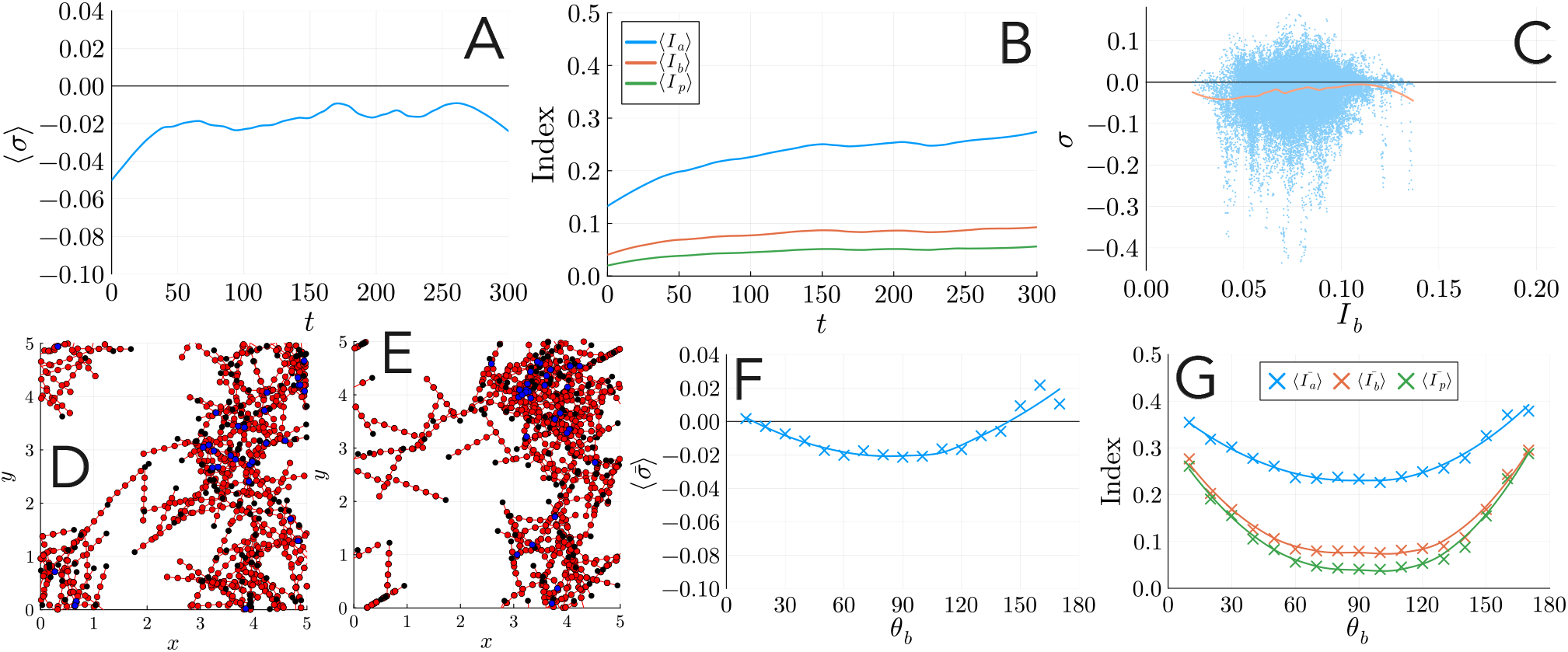
Numerical results for simulations with 70° branching turnover with rate *k*_off,*a*_ = 0.04 s^*−*1^. (A) Trial-averaged mean normal stress over time for *N* = 10 simulations, smoothed using a Savitzky–Golay filter (LOESS regression). (B) Trial-averaged bundle indices over time for *N* = 10 simulations, smoothed using a Savitzky–Golay filter (LOESS regression). (C) Comparison of instantaneous measurements (blue dots) of mean normal stress and bundle index across all simulations and time. The solid curve is a LOESS regression of the data. (D) Example network configuration at *t* = 50. (E) Configuration of the same network at *t* = 300. (F) The effect of branching angle on time-averaged and trial-averaged stress. (G) The effect of branching angle on time-averaged and trial-averaged bundle indices. We performed *N* = 10 simulations for each value of *θ*_*b*_ (crosses), and smoothed using a Savitzky–Golay filter (solid curves).

As Figure 7G–H show, the angle *θ*_*b*_ = 70° is close to the optimal angle for maximising contractility and decreasing aggregation, bundle formation, and parallel-bundle formation compared to other potential branching angles. Therefore, 70° branching appears to be a non-uniform turnover mechanism that disrupts bundles, favouring contractility over bundle formation. This finding reflects the results of Ennomani et al. [31], which experimentally support the higher contractility of branched networks.

### 3.5 Treadmilling promotes bundling while maintaining contractility

Unlike uniform turnover, biased turnover, and branching, treadmilling does not involve the removal and replacement of filaments in the network. Instead, existing filaments lose actin from their minus-ends, gain actin at their plus-ends, but otherwise maintain their position. Figure 8A shows that treadmilling prevents the loss of contractility that occurs with no turnover. Instead, the trend in contractility is similar to other non-uniform turnover methods, featuring a loss of contractility over the first approximately 50 s, before settling in a weakly-contractile state. However, unlike biased turnover and branching, with treadmilling Figure 8B shows that aggregation, bundle formation, and parallel-bundle formation continue to increase over a longer time. The bundle-formation index eventually reaches a similar value to that attained with no turnover, suggesting that treadmilling must cause the persistent contractility observed in Figure 8A. The results in Figure 8C, where weaker contractility remains possible with larger values of *I*_*b*_, reinforce this idea. Treadmilling lessens the negative feedback between bundle formation on contractility, enabling filaments within these bundles to retain net contractility.

**Figure 8:**
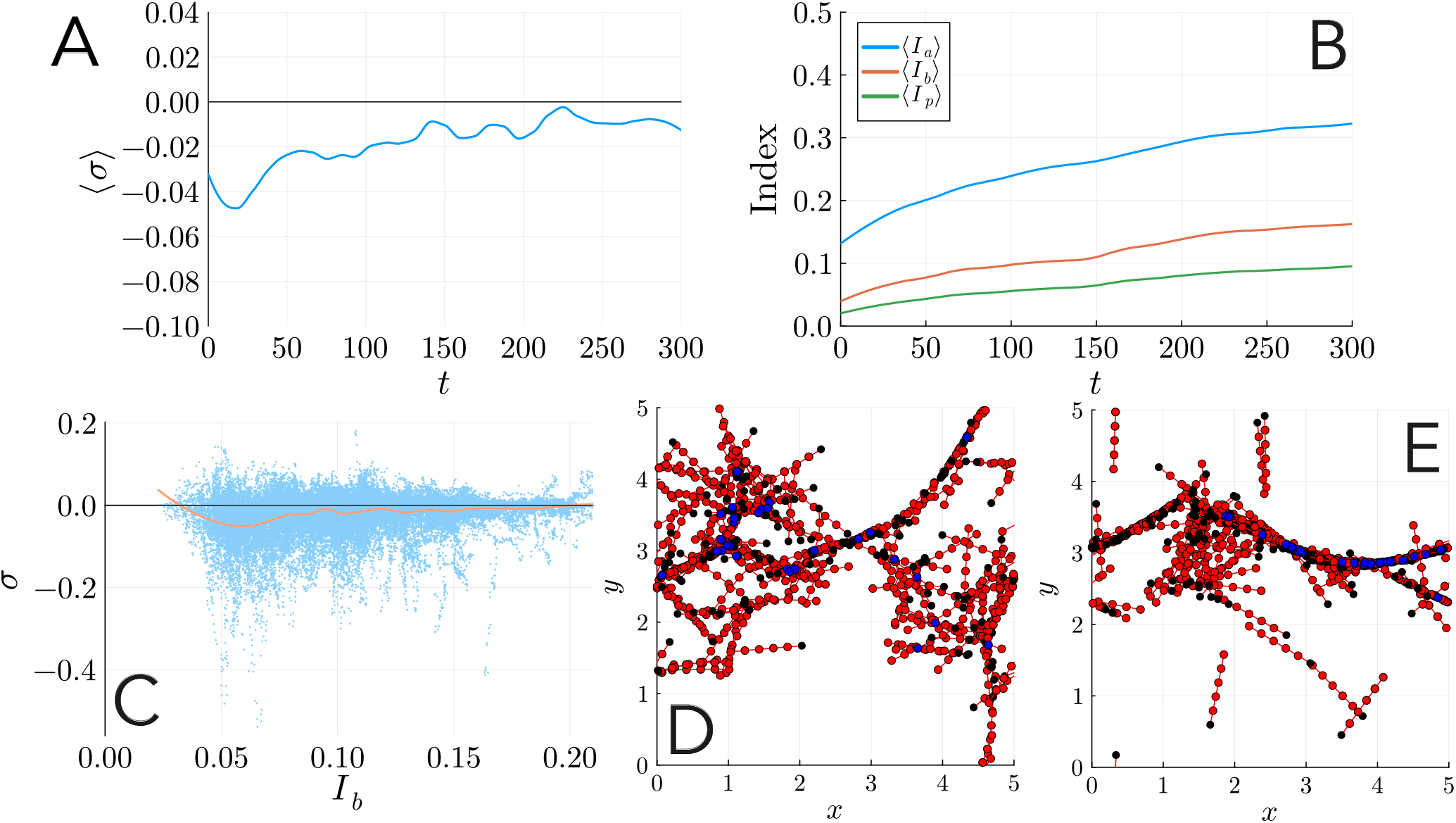
Numerical results for simulations with treadmilling turnover with rate *k*_off,*a*_ = 0.04 s^*−*1^. (A) Trial-averaged mean normal stress over time for *N* = 10 simulations, smoothed using a Savitzky–Golay filter (LOESS regression). (B) Trial-averaged bundle indices over time for *N* = 10 simulations, smoothed using a Savitzky–Golay filter (LOESS regression). (C) Comparison of instantaneous measurements (blue dots) of mean normal stress and bundle index across all simulations and time. The solid curve is a LOESS regression of the data. (D) Example network configuration at *t* = 50. (E) Configuration of the same network at *t* = 300.

The network configurations formed with treadmilling are also distinct from those formed with other turnover mechanisms. In Figure 8D, a long, thin curved filament bundle reminiscent of an unattached stress fibre emerges in the upper part of the domain. Eventually, most filaments become part of a bundle-like aggregate, as shown in Figure 8E. Since most of the filament maintains its position during treadmilling, these patterns are similar to the network patterns with no turnover, in Figure 4F. Unlike with no turnover, these networks with treadmilling maintain contractility. Minus-end disassembly provides a possible qualitative explanation for this persistent contractility [12]. As the motors move towards filament plus-ends, the minus-ends are pushed apart, which could lead to expansion [34]. The minus-end disassembly diminishes this expansive potential, resulting in net contractility. Treadmilling may also cause the plus-ends and minus-ends locations to more closely resemble the periodic arrangement in contractile stress fibres [41, 42].

### 3.6 Filament bending enhances bundle formation and protein friction enhances contractility for non-uniform turnover

We conclude our investigation by exploring how mechanical parameters affect contractility and bundle formation for each turnover model. The most important parameters were flexural rigidity and the protein-friction coefficient, and Figure 9 summarises their effects. The background-drag coefficient and reference motor off-rate were less important (see the Supplementary Material for details), and we assumed the spring constants, free-moving motor velocity and stall force to be fixed based on experimental data. These results accord with our previous study [28], where filament flexibility (reduced flexural rigidity) and protein friction promoted contractility.

**Figure 9:**
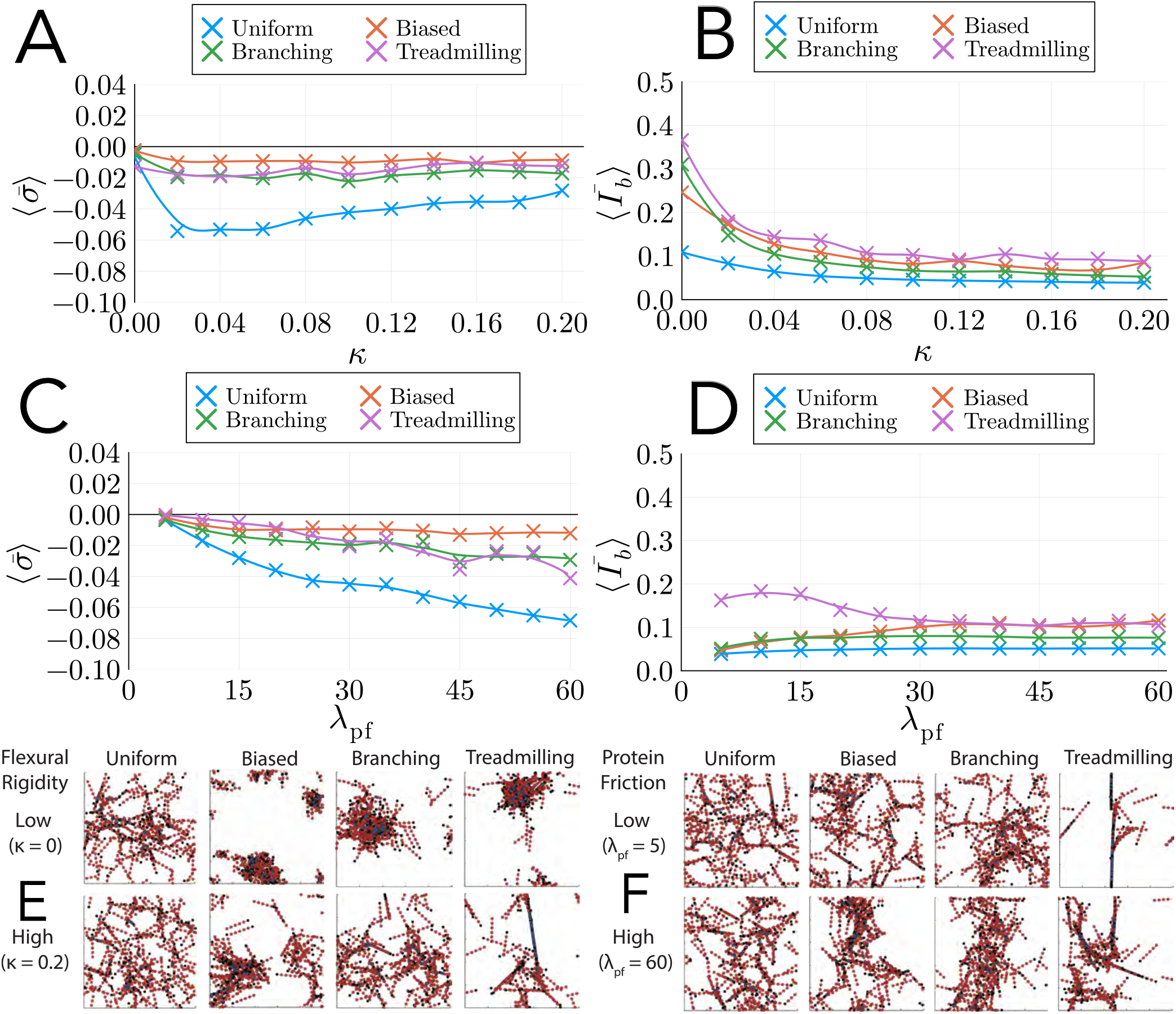
Numerical results for simulations with different forms of turnover with rate *k*_off,*a*_ = 0.04 s^*−*1^, and varying flexural rigidity *κ* and protein friction coefficient *λ*_pf_. (A) The effect of *κ* on trial-averaged mean normal stress over time for *N* = 10 simulations (crosses). Solid curves are obtained by applying a Savitzky–Golay filter (LOESS regression). (B) The effect of *κ* on bundle formation index over time for *N* = 10 simulations (crosses). Solid curves are obtained by applying a Savitzky–Golay filter (LOESS regression). (C) The effect of *λ*_pf_ on trial-averaged mean normal stress over time for *N* = 10 simulations (crosses). Solid curves are obtained by applying a Savitzky–Golay filter (LOESS regression). (D) The effect of *λ*_pf_ on trial-averaged bundle formation index over time for *N* = 10 simulations (crosses). Solid curves are obtained by applying a Savitzky–Golay filter (LOESS regression). (E) The effect of flexural rigidity on network configuration for different turnover models. (F) The effect of protein-friction coefficient on network configuration for different turnover models.

Low flexural rigidity *κ* decreases resistance to filament bending. With uniform turnover, decreased *κ* increases network contractility until *κ* = 0.02 pN µm^2^, as Figure 9A shows. However, flexural rigidity has less impact on contractility for all methods of non-uniform turnover. Decreasing *κ* increases bundle formation (Figure 9B), because flexible filaments can more readily remodel into an aggregate due to myosin activity (see Figure 9E). These aggregates are less contractile than a disordered network due to the trade-off between bundle formation and contractility. Uniform turnover disrupts the aggregates and helps to maintain network contractility. In contrast, with non-uniform turnover the aggregates persist, because new filaments are more likely to emerge within the aggregate itself. As *κ* increases, branching and treadmilling disperse the aggregate more than biased turnover (see Figure 9E). Consequently, networks with branching and treadmilling yield networks that are more contractile compared to networks with biased turnover. However, although branching and treadmilling promote contractility more effective than biased turnover, they are less effective than uniform turnover regardless of *κ*.

Increasing the protein-friction coefficient, *λ*_pf_, increases contractility with all forms of turnover, as Figure 9C shows. Increasing *λ*_pf_ increases bundle formation for biased turnover and branching, as Figure 9D and the network snapshots shown in Figure 9F indicate. Since increasing *λ*_pf_ elevates both contractility and bundle formation with biased turnover and branching, strong protein friction (for example due to cross-linking) may be important for generating persistent contractile bundles. With uniform turnover, *I*_*b*_ to be largely independent of *λ*_pf_ because uniform turnover effectively disrupt bundles regardless of *λ*_pf_. With uniform turnover, there is large increase in contractility as *λ*_pf_ increases. In contrast, with biased turnover and branching, *I*_*b*_ increases as *λ*_pf_ increases. Due to the trade-off between bundle formation and contractility, this results in a smaller gain in contractility for these turnover methods as *λ*_pf_ increases, compared to uniform turnover.

For treadmilling turnover, increasing *λ*_pf_ decreases bundle formation, which is the opposite behaviour to biased turnover and branching (see Figure 9B). When *λ*_pf_ = 5 pN µm^*−*1^ s (weak protein friction), networks with treadmilling form a very thin bundle (see Figure 9F) with high *I*_*b*_. Small *λ*_pf_ corresponds to reduced resistance to relative motion between overlapping filaments, allowing the network to more easily remodel into the thin bundle structure, which resembles the stationary patterns produced with no turnover. With treadmilling, this pattern remains only weakly contractile. For low values of *λ*_pf_, treadmilling yields less contractility than biased turnover or branching, due to the large discrepancy in *I*_*b*_. As *λ*_pf_ increases, relative motion of overlapping filaments becomes more difficult, and the network with treadmilling remodels more slowly. For *λ*_pf_ = 60 pN µm^*−*1^ s, Figure 9F shows that the network no longer forms a single thin bundle. For treadmilling, this causes *I*_*b*_ to decreases as *λ*_pf_ increases. Contractility then increases with increasing *λ*_pf_, due to both decreased bundle formation, and the general contractile effect of protein friction. For large values of *λ*_pf_, the treadmilling yields increased contractility compared to biased turnover and branching. This increased contractility occurs despite the treadmilling networks maintaining *I*_*b*_ values similar to biased turnover networks for large *λ*_pf_. The increased contractility with high *I*_*b*_ further reinforces the idea that treadmilling lessens the trade-off between bundle formation and contractility.

## 4 Discussion and Conclusion

Filament turnover is a key mediator of actomyosin-network architecture and contractility. In this work, we implemented an agent-based model of an actomyosin network with four turnover models: uniform, biased, branching, and treadmilling. These turnover models are simplified representations of the complex molecular processes involved in network remodelling. We used the model to investigate the interplay between contractility, bundle formation, and turnover. Figure 10 provides an overview of our results. Without turnover, there is negative feedback between bundle formation and contractility. Disordered networks initially contract, forming bundles. These bundles feature filaments aligned in parallel, similar to the aster formation in previous studies [11, 28, 58, 59]. Eventually, these parallel structures become stationary, preventing further contractility. Uniform turnover, branching, and treadmilling can prolong contractility. Introducing uniform turnover or branching disrupts bundle formation, allowing network contractility to persist, whereas treadmilling disrupts the trade-off between bundle formation and continued contractility. In contrast, biased turnover increases bundle formation, which quickly results in reduced contractility. Mechanical factors also play a role in maintaining contractility and bundle formation. In our simulations, the clearest mechanical effects were that increased filament bending and protein friction promote contractility. Taken together, these results indicate that a combination of mechanically-driven contractility and bundle-disrupting turnover is necessary to sustain network contractility and elevated bundle formation in initially-disordered networks.

**Figure 10:**
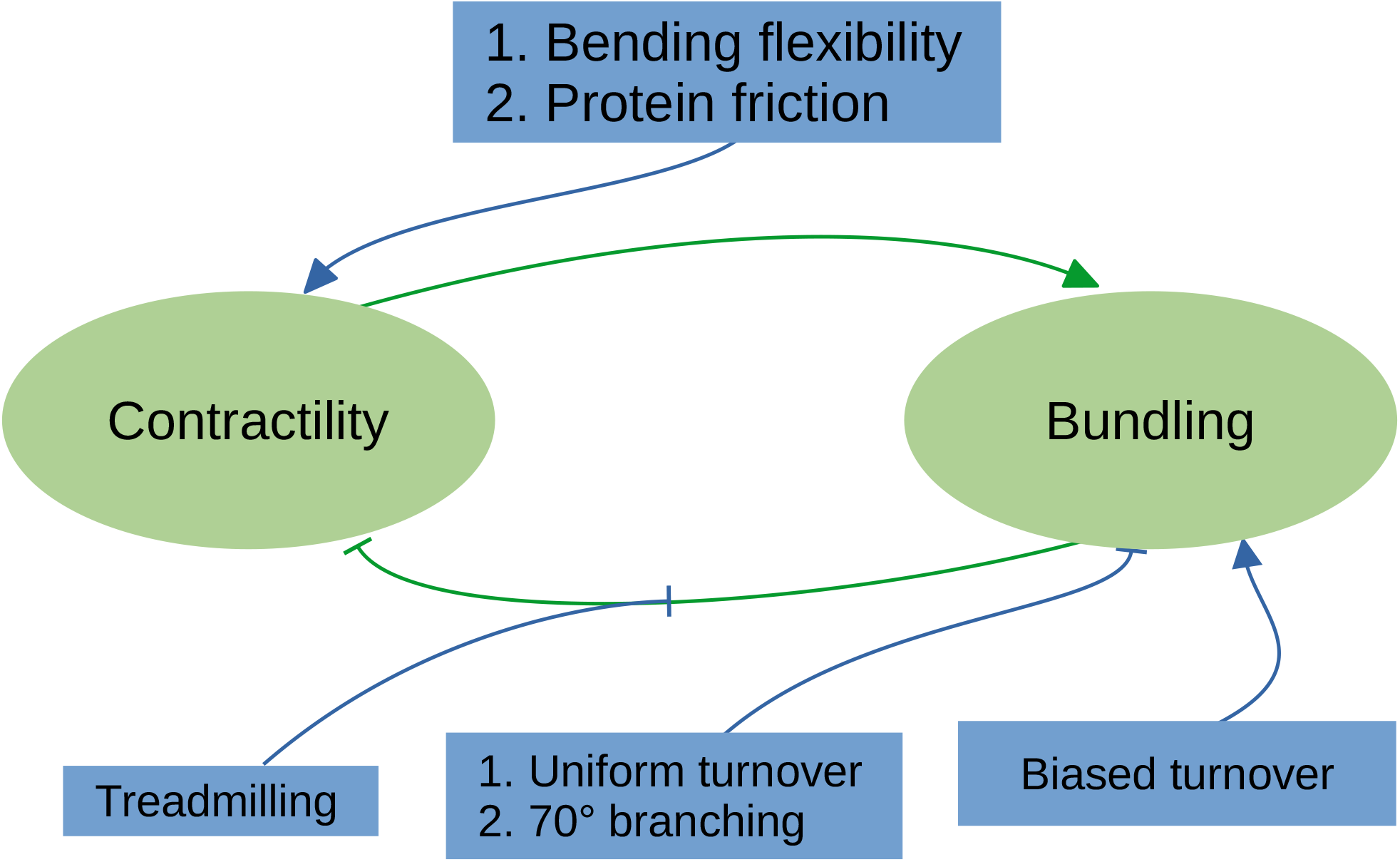
The interplay between contractility and bundling (green ellipses), and turnover methods and mechanical factors (blue rectangles) in initially-disordered actomyosin networks. Lines between each group indicate the direction of causality. Arrow heads indicate factors that positively influence each other, and perpendicular lines indicate factors that negatively influence each other.

In this study, we focused on isolating the effects of individual turnover models working alone. A realistic model for filament turnover in the cell might involve a combination of the turnover models that we investigated. Our work suggests that some individual mechanisms have opposing effects on the trade-off between contractility and bundle formation. For example, uniform turnover and branching disrupt bundles and promotes contractility, whereas biased turnover promotes bundle formation, hastening loss of contractility. A combination of turnover methods working in concert, might allow cells to tune the levels of contractility and bundle formation, depending on the scenario. The rate of each turnover method might also influence this optimal combination. Our general findings on the mechanisms of turnover do not rely on specific turnover rates. However, we also present quantitative details on how turnover rate, *k*_off,a_, impacts contractility and bundle formation in the Supplementary Material.

Some drawbacks of our approach need to be considered when interpreting the results. Since the cell cortex is a thin layer, for simplicity we used a 2D model. Extending the model to 3D would require explicit simulation of motor and cross-linker binding and unbinding, because the assumption that motors bind at intersections in 2D is unavailable in 3D. Modelling freely-moving populations of motors and cross-linker proteins in the cytoplasm might influence bundle formation by localising motor activity, a possibility our model neglects. We also cannot rule out the possibility that domain and network size effects might impact our results, although we obtained similar qualitative results for smaller simulations with 75 filaments and 15 motors on 2.5 µm *×* 2.5 µm domains (results not shown). The computational cost of the simulations constrains our ability to verify the results on larger scales. Another weakness is that we do not compute localised forces and stress throughout the domain, but instead adopt a simplified model based on uniform deformation. Accounting for non-uniform directed and localised forces is a difficult problem beyond the scope of this study [92]. Our objective is to compare modelling and turnover scenarios, rather than accurately modelling the active material mechanics.

The filament turnover and mechanics investigated here might be relevant to the contractile ring and early stress-fibre formation. In the cell, focal-adhesion sites attach filament bundles to the cytoskeleton. Although Vignaud et al. [14] showed that unattached stress fibres transmit tension due to their connection with the surrounding network, focal adhesions help stress fibres transmit forces through the cell [15, 93]. Furthermore, initially-unattached filament bundles can eventually attach to focal adhesion sites [16, 94, 95]. We only consider bundles that are not attached to focal adhesions. However, it is more common for stress fibres to be attached to focal adhesions than not attached [96]. An extension for future work could be to introduce focal adhesions into the simulations, modelling them as isolated regions of large background drag. A realistic view of biological networks might also feature multiple turnover models acting simultaneously. Combining uniform and non-uniform turnover might allow networks to tune contractility and bundle formation, providing another avenue for future extension.

## Supporting material

An electronic supplement containing additional simulations results is available with the article.

## Author contributions

- **AKYT:** Conceptualization, methodology, software, formal analysis, investigation, data curation, writing — original draft, visualization.
- **AM:** Conceptualization, writing — review & editing, supervision, funding acquisition.
- **DBO:** Conceptualization, methodology, formal analysis, writing — review & editing, supervision, funding acquisition.

## Acknowledgements

We acknowledge funding from the Australian Research Council (Grant numbers DP180102956, DP230100406, and DE240100097). We also acknowledge Claudia Blom, whose summer research project informed the spatial statistics used in this work.

