## Supporting material for "Non-uniform filament turnover and mechanically-driven contractility and bundle formation in disordered actomyosin networks"

#### A Default Parameters

The default parameters used in simulations and their sources are shown in Table A.1.

Table A.1: Default parameters for the actomyosin network simulations.

| Parameter | Symbol | Value | Units | Source |
|---|---|---|---|---|
| Filament spring stiffness | $k_a$ | 1000 | $\mathrm{pN\,\mu m^{-1}}$ | [1] |
| Filament–background drag coefficient | $\lambda_a$ | 0.05 | $\mathrm{pN\,\mu m^{-2}\,s}$ | [2–4] |
| Filament flexural rigidity | $\kappa$ | 0.073 | $\mathrm{pN\,\mu m^{2}}$ | [5] |
| Protein-friction drag coefficient | $\lambda_{\mathrm{pf}}$ | 30 | $\mathrm{pN\,\mu m^{-1}\,s}$ | [6] |
| Filament-turnover rate | $k_{\mathrm{off},a}$ | 0.04 | $\mathrm{s^{-1}}$ | [7, 8] |
| Motor spring constant | $k_m$ | 1000 | $\mathrm{pN\,\mu m^{-1}}$ | [1] |
| Motor stall force | $F_s$ | 5 | pN | [9–11] |
| Load-free motor velocity | $V_m$ | 0.5 | $\mathrm{\mu m\,s^{-1}}$ | [9, 11, 12] |
| Motor reference off-rate | $k_{\mathrm{off},m}$ | 0.35 | $\mathrm{s^{-1}}$ | [13, 14] |
| Reference force for motor unbinding | $F_{\mathrm{ref}}$ | 12.6 | pN | [15] |
| Filament branching angle | $\theta_b$ | $70^\circ$ | — | [16] |
| Standard deviation for translations | $\sigma_x$ | 0.05 | — | Assumption |
| Standard deviation for rotations | $\sigma_\theta$ | $\pi/20$ | — | Assumption |

### B Supplementary Results and Discussion

This appendix contains more results from the simulations to support the main text. We explore how turnover rate impacts contractility and bundle formation for each turnover model. We also explore how the polarity-reversal probability affects biased turnover, based on three main scenarios: same polarity, mixed polarity, and reversed polarity. Same-polarity biased turnover refers to all new filaments having the same polarity as the reference filament, and promotes parallel-bundle formation. Mixed polarity is the scenario considered in the main manuscript, where new filaments have the same polarity as the reference filament with probability 0.5. Reversed polarity involves new filaments having the opposite polarity to the reference filament. We also compare simulations with mechanics versus simulations with turnover only, isolating the key contribution of filament and motor mechanics to bundle formation. Finally, we present results obtained by varying the background-drag coefficient and the motor-detachment rate. Compared to flexural rigidity and protein friction, these parameters do not significantly impact contractility or bundle formation.

#### B.1 Trade-off between bundle formation and contractility

To confirm the trade-off between bundle formation and contractility, we simulated 10 networks and computed the mean normal stress $\sigma$ and bundle formation indices $I_a$, $I_b$, and $I_p$ at each time step in the simulations. Figure B.1 illustrates a smoothed mean of the 10 simulations, providing representative illustration of how stress and bundle formation develop over time. We use the same bar and angle-bracket notation for averaged quantities as the main manuscript.

As Figure B.1a shows, for $k_{\text{off},a} = 0.04\,\text{s}^{-1}$ contractility persists with uniform turnover, but contractility decreases with time for non-uniform (biased, branching, and treadmilling) turnover. Although faster turnover enables contractility to persist longer for branching and treadmilling turnover (see Figure B.1e), uniform turnover produces the strongest contractility. Uniform turnover increases the randomness in filament positions compared to biased turnover, where the position of new filaments depends on the positions of existing filaments. This prevents formation of stationary structures, enabling persistent contractility. However, as Figure B.1b shows, the random network remodelling of uniform turnover reduces bundle formation. In contrast, all

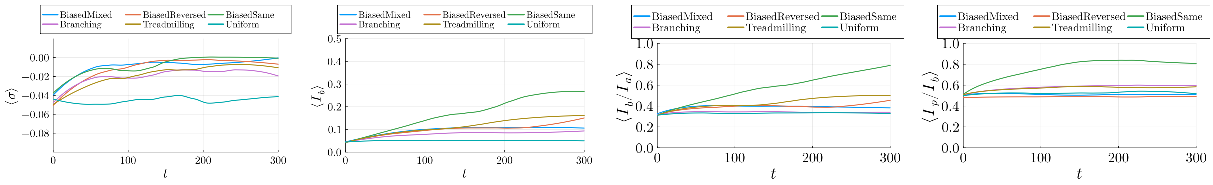


(a) $k_{\text{off},a} = 0.04\,\text{s}^{-1}$. (b) $k_{\text{off},a} = 0.04\,\text{s}^{-1}$. (c) $k_{\text{off},a} = 0.04\,\text{s}^{-1}$. (d) $k_{\text{off},a} = 0.04\,\text{s}^{-1}$.

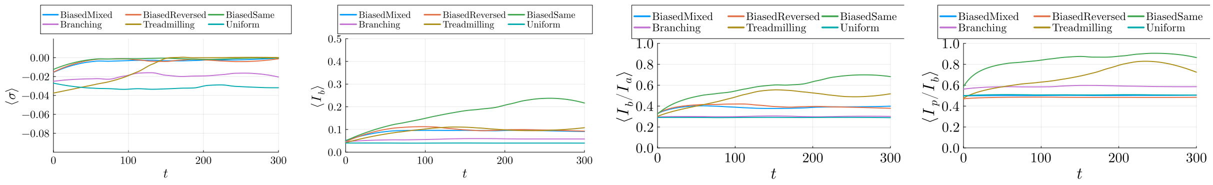


(e) $k_{\text{off},a} = 0.2\,\text{s}^{-1}$. (f) $k_{\text{off},a} = 0.2\,\text{s}^{-1}$. (g) $k_{\text{off},a} = 0.2\,\text{s}^{-1}$. (h) $k_{\text{off},a} = 0.2\,\text{s}^{-1}$.

Figure B.1: Smoothed trial-averaged ($N = 10$) results for how mean trial-averaged normal stress and bundle formation indices evolve over time, for different models of turnover. (a) Mean normal stress $\langle\sigma\rangle$, for turnover rate $k_{\text{off},a} = 0.04\,\text{s}^{-1}$. (b) Bundle formation index, $\langle I_b\rangle$, for $k_{\text{off},a} = 0.04\,\text{s}^{-1}$. (c) Bundle-aggregate ratio, $\langle I_b/I_a\rangle$, for $k_{\text{off},a} = 0.04\,\text{s}^{-1}$. (d) Parallel bundle ratio, $\langle I_p/I_b\rangle$, for $k_{\text{off},a} = 0.04\,\text{s}^{-1}$. (e) Mean normal stress $\langle\sigma\rangle$, for $k_{\text{off},a} = 0.2\,\text{s}^{-1}$. (f) Bundle formation index, $\langle I_b\rangle$, for $k_{\text{off},a} = 0.2\,\text{s}^{-1}$. (g) Bundle-aggregate ratio, $\langle I_b/I_a\rangle$, for $k_{\text{off},a} = 0.2\,\text{s}^{-1}$. (h) Parallel bundle ratio, $\langle I_p/I_b\rangle$, for $k_{\text{off},a} = 0.2\,\text{s}^{-1}$.

non-uniform turnover methods cause increased bundle formation over time compared to a random initial condition.

To further quantify the simulation results, we consider the bundle-aggregate ratio, $I_b/I_a$, and parallel-bundle ratio $I_p/I_b$. The bundle-aggregate ratio indicates whether bundle formation is increasing relative to general aggregation, and the parallel-bundle ratio indicates the relative bundle orientations. The random initial condition corresponds to approximately $I_b/I_a = 0.19$ and $I_p/I_b \approx 0.5$. In Figures B.1c, B.1d, B.1g and B.1h, increases in $I_b$ accompany commensurate rises in bundle-aggregate ratio and parallel-bundle ratio. This suggests that increased bundling is explained by formation of parallel structures, which are less contractile than antiparallel structures. Taken together, the results in Figure B.1 show the trade-off between persistent contractility and bundle formation in actomyosin networks that others have previously established [17–19].

Figure B.2 further illustrates the trade-off between contractility and bundle formation. These plots display time-averaged mean normal stress, $\bar{\sigma}$, and bundle indices, versus turnover rate $k_{\text{off},a}$, averaged over $N = 10$ simulations for each value of $k_{\text{off},a}$. Figure B.2a shows maximum time-averaged contractility with uniform turnover. Uniform turnover maximises time-averaged contractility because contractility persists with uniform turnover, but does not with non-uniform turnover. However, simulations with uniform turnover show less aggregation, bundle formation, and parallel-bundle formation than simulations with non-uniform turnover. Bundle formation tends to decrease with increasing turnover rate, as faster turnover disperses bundles and prevents stationary bundle formation.

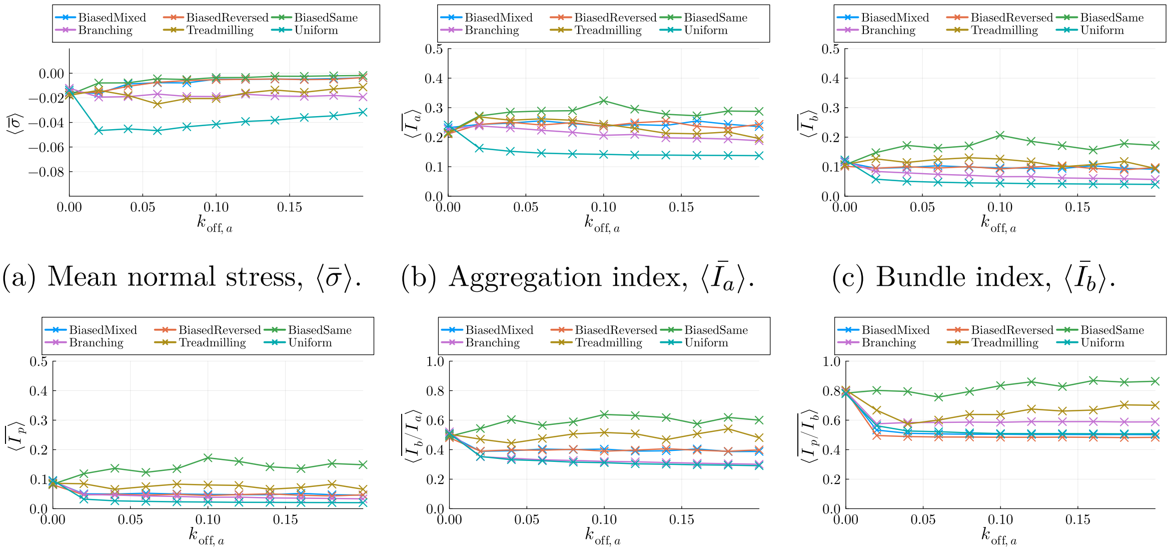


(a) Mean normal stress, $\langle\bar{\sigma}\rangle$. (b) Aggregation index, $\langle\bar{I}_a\rangle$. (c) Bundle index, $\langle\bar{I}_b\rangle$.

(d) Parallel index, $\langle\bar{I}_p\rangle$. (e) Bundle ratio, $\langle\overline{I_b/I_a}\rangle$. (f) Parallel ratio, $\langle\overline{I_p/I_b}\rangle$.

Figure B.2: Trial-averaged and time-averaged mean normal stress, $\langle\bar{\sigma}\rangle$, and bundle formation index, $\langle\bar{I}_b\rangle$, versus turnover rate, $k_{\mathrm{off},a}$, for six different methods of turnover. Presented results are the mean of $N = 10$ simulations for each value of the turnover rate.

#### B.2 Biased turnover increases bundle formation and leads quickly to loss of contractility

A clear result from Figures B.1 and B.2 is that biased turnover causes loss of contractility over time, regardless of whether the polarity of new filaments is the same, mixed, or reversed. Same polarity biased turnover also yielded large increases in aggregation, bundle formation, and parallel-bundle formation. In contrast, mixed and reversed polarity biased turnover yielded more bundle formation relative to aggregation than uniform turnover (see Figures B.1c and B.1g), but negligible change to the parallel-bundle ratio compared to uniform turnover (see Figures B.1d and B.1h). To investigate these effects in more detail, we computed solutions using biased turnover, but varying polarity-reversal probability, $P_{\mathrm{rev}}$, and standard deviations $\sigma_x$ and $\sigma_\theta$. Figures B.3a–B.3d show that polarity reversal probability has little effect on contractility and bundle formation, except when $P_{\mathrm{rev}} = 0$. As expected, if $P_{\mathrm{rev}} = 0$ increases in overall bundle formation are explained by increases in parallel-bundle formation.

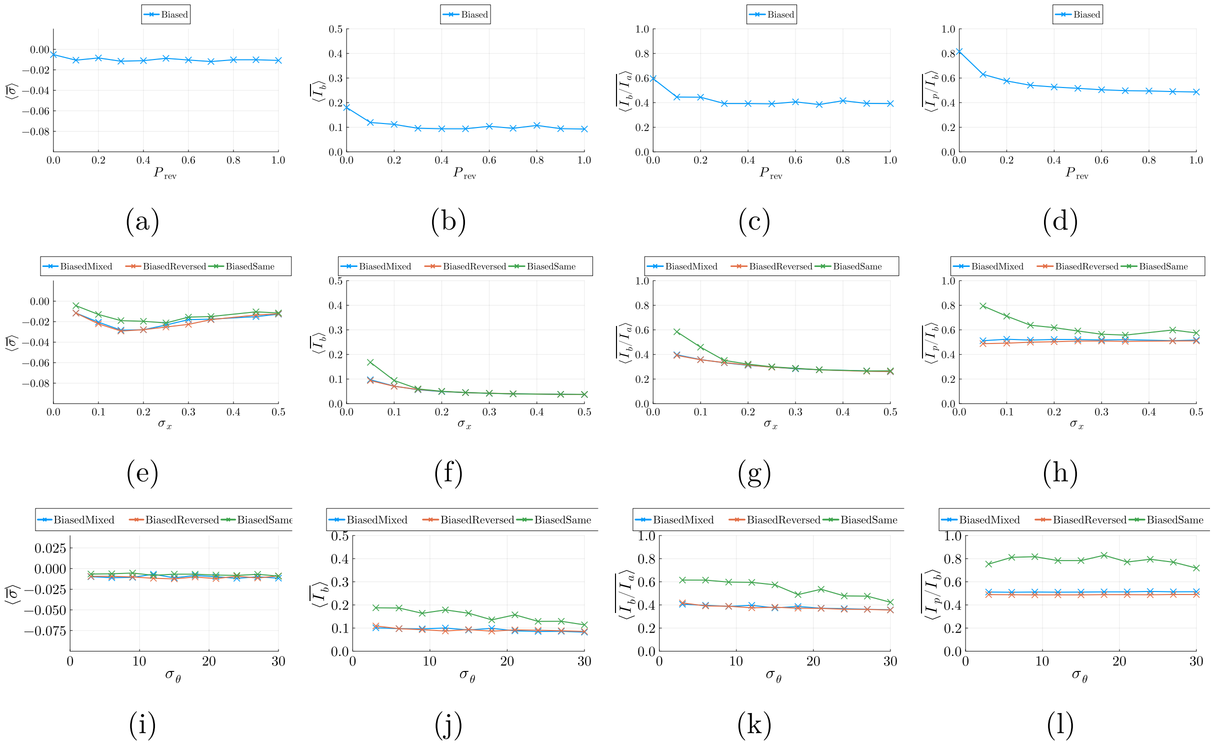


Figure B.3: The effect of polarity reversal probability, $P_{\text{rev}}$ and standard deviation for translations, $\sigma_x = \sigma_y$, on contractility and bundle formation with biased non-uniform turnover.

Results with varying $\sigma_x$, the standard deviation used when translating new filaments, reveals similar trends. Increases to $\sigma_x$ cause new filaments to be formed further from the reference filament, thereby decreasing the bundle index and parallel-bundle ratio, especially for same-polarity biased turnover. Figure B.3e suggests that $\sigma_x \approx 0.15$ is optimal for generating contractility. For smaller values $\sigma_x < 0.15$, new filaments may tend to overlap existing ones, favouring bundle formation but not continued contractility. In contrast, $\sigma_x$ too large may limit network connectivity, decreasing contractility. For $\sigma_x \geq 0.15$ most of the additional bundle formation due to biased turnover is lost, providing another illustration of the trade-off between bundle formation and contractility. The standard deviation for the angle of rotation, $\sigma_\theta$, appears to have little effect on contractility, but larger angles decrease bundle formation and parallel-bundle formation, as expected.

#### B.3 70° branching favours contractility over bundle formation

We now focus on the effect of the branching angle in branching turnover. Figure B.2c shows that branching enhances bundle formation compared to simulations with uniform turnover, for all turnover rates $k_{\text{off,a}}$ tested. However, although with branching turnover contractility persists over time, Figure B.2c indicates that branching decreases overall contractility compared to uniform turnover. We investigate whether a $\theta_b = 70°$ branching angle maximises contractility compared to other possible angles. To achieve this, we simulate networks with fixed branching angle (10 simulations for each value of $\theta_b$), while quantifying contractility and bundle formation. We also illustrate the effect of filament and motor mechanics, by comparing results to solutions with no mechanics (turnover only, no motor movement).

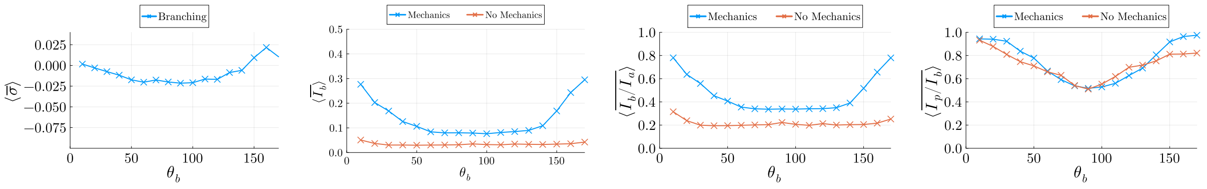


(a) (b) (c) (d)

Figure B.4: Trial-averaged and time-averaged mean normal stress, $\langle\bar{\sigma}\rangle$, bundle-formation index, $\langle\bar{I}_b\rangle$, bundle ratio, $\langle \overline{I_b/I_a}\rangle$, and parallel-bundle ratio, $\langle \overline{I_p/I_b}\rangle$, versus branching angle, $\theta_b$.

Figure B.4a illustrates the impact of branching angle, $\theta_b$, on contractility. In simulation with mechanics, branching turnover gave rise to network contractility for all values of $\theta_b$ between $\theta_b = 20°$ and $\theta_b = 140°$. Our simulations show that branching with angles between $\theta_b = 60°$ and $\theta_b = 100°$ is optimal for generating contraction. Small values of $\theta_b$ produce new filaments that are closely aligned to existing filaments, which only enables a small amount of contractility. In contrast, values of $\theta_b$ close to 180° produce new filaments aligned similarly to existing filaments, but with reversed polarity. As Figure B.4a shows, branching turnover with $\theta_b \geq 150°$ gives rise to expansion, rather than contraction. Interestingly, these results contradict the contractile effect of reversed-polarity biased turnover, which also generates initially antiparallel assemblies. A possible explanation is the role of F-actin bending in contractility [20]. Depending on the relative filament positions, reversed-polarity biased turnover often produces motor-filament assemblies whose expansive effect is not inhibited by bending. Compared to other possible branching angles, branching with $\theta_b = 70°$ favours contractility, rather than bundle formation.

As Figure B.4b shows, smaller branching angles increase bundle formation significantly compared to branching at other angles. The increase in $I_b$ at these angles occurs because the new filaments are oriented similarly to existing filaments, within the 20° threshold to be included in the bundle index. However, for other values of $\theta_b$ the new filaments are not aligned similarly to existing ones. The inclusion of mechanics increases bundle formation for all branching angles $\theta_b$, compared to results with branching turnover only, and no mechanics. Furthermore, Figures B.4c and B.4d suggest that mechanics generates a large increase in bundle formation relative to aggregation, but only a small increase in parallel-bundle formation. With branching, filament and motor mechanics accelerate generation of bundles from aggregates, which we now explore in further detail.

#### B.4 Filament bending enhances bundle formation and protein friction enhances contractility for non-uniform turnover

Appendix B.1 explored the effect of varying the rate and method of turnover, holding mechanical parameters constant. We now investigate how filament and motor mechanics affect bundle formation compared to simulations with no mechanics (turnover only, with no filament or motor movement). Since networks without mechanics generate no contraction, Figure B.2A confirms that filament and motor mechanics enhance contractility for all turnover rates methods tested. Filament and motor mechanics also enhance bundle formation for each turnover rate and method, as Figure B.5 shows. Uniform and branching turnover, filament and motor mechanics increase both overall bundle formation and bundle formation relative to aggregation.

At low turnover rates, the increase in bundle formation accompanies an increase in parallel-bundle formation. For faster turnover rates, uniform and branching turnover give increased bundle formation without an increase in parallel-bundle formation. Although less pronounced, a similar result emerges with biased turnover with mixed or reversed polarity. For all turnover rates tested, adding mechanics yields large increases in bundle formation and bundles-aggregate ratio. With same-polarity biased turnover, this increase in bundle formation is explained by formation of parallel bundles. Parallel-bundle formation is less pronounced for treadmilling turnover, especially with intermediate turnover rates. Interestingly, $k_{\mathrm{off},a} = 0.04\,\mathrm{s}^{-1}$, minimises mechanically-driven parallel-bundle formation with treadmilling turnover. Overall, these results suggest a key role for mechanics in generating bundles that might remain contractile, especially with branching and treadmilling turnover.

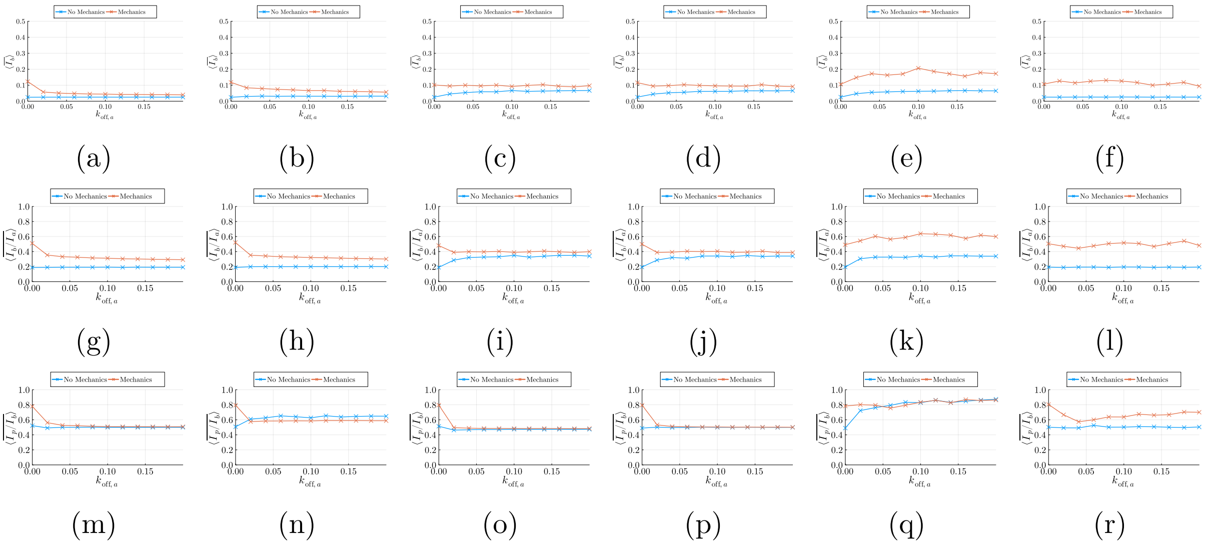

(a) (b) (c) (d) (e) (f)

(g) (h) (i) (j) (k) (l)

(m) (n) (o) (p) (q) (r)

Figure B.5: Trial-averaged and time-averaged bundle formation index, $\langle \bar{I}_b \rangle$, versus turnover rate, $k_{\text{off},a}$, with and without mechanics. Presented results are the mean of $N = 10$ simulations for each value of the turnover rate. (a, g, m) Uniform turnover. (b, h, n) Branching turnover. (c, i, o) Biased turnover (reversed polarity). (d, j, p) Biased turnover (mixed polarity). (e, k, q) Biased turnover (same polarity). (f, l, r) Treadmilling.

#### B.5 The background-drag coefficient and the motor-detachment rate have little effect on contractility and bundle formation

The results in the main manuscript show how the filament flexural rigidity and the protein-friction coefficient impact contractility and bundle formation. Figure B.6 illustrates how contractility and bundle formation evolve over time, complementing the time-averaged results presented in the main text. The results reflect similar trends for each turnover model as Figure B.1.

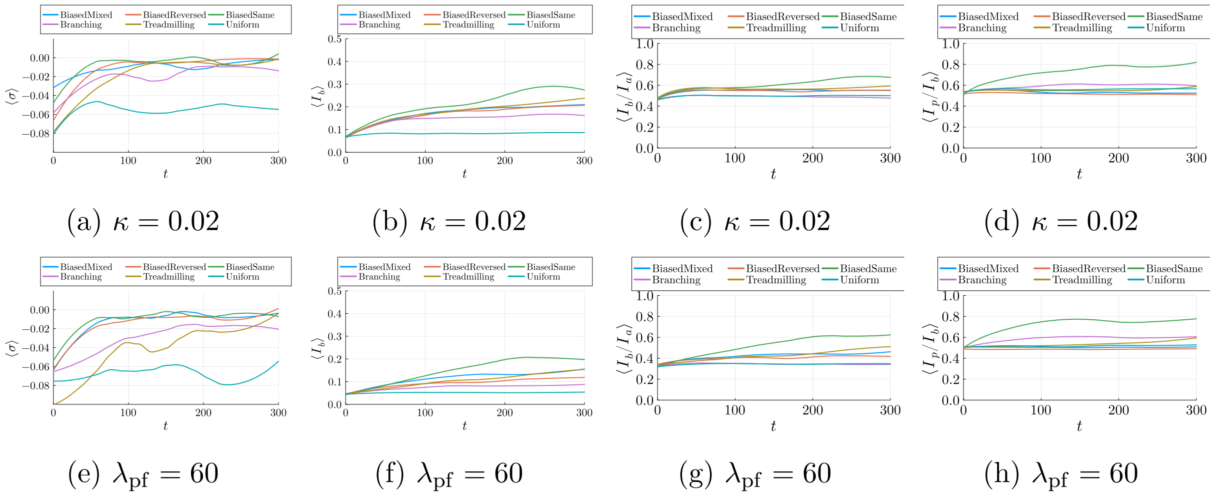


(a) $\kappa = 0.02$ (b) $\kappa = 0.02$ (c) $\kappa = 0.02$ (d) $\kappa = 0.02$

(e) $\lambda_{\text{pf}} = 60$ (f) $\lambda_{\text{pf}} = 60$ (g) $\lambda_{\text{pf}} = 60$ (h) $\lambda_{\text{pf}} = 60$

Figure B.6: The impacts of low flexural rigidity and high protein friction on time-dependent contractility and bundle formation, with $k_{\text{off},a} = 0.04$ and different turnover models.

Compared to filament flexural rigidity and the protein-friction coefficient, varying the background-drag coefficient and the motor-detachment rate does not significantly affect contractility and bundle formation, as Figure B.7 indicates.

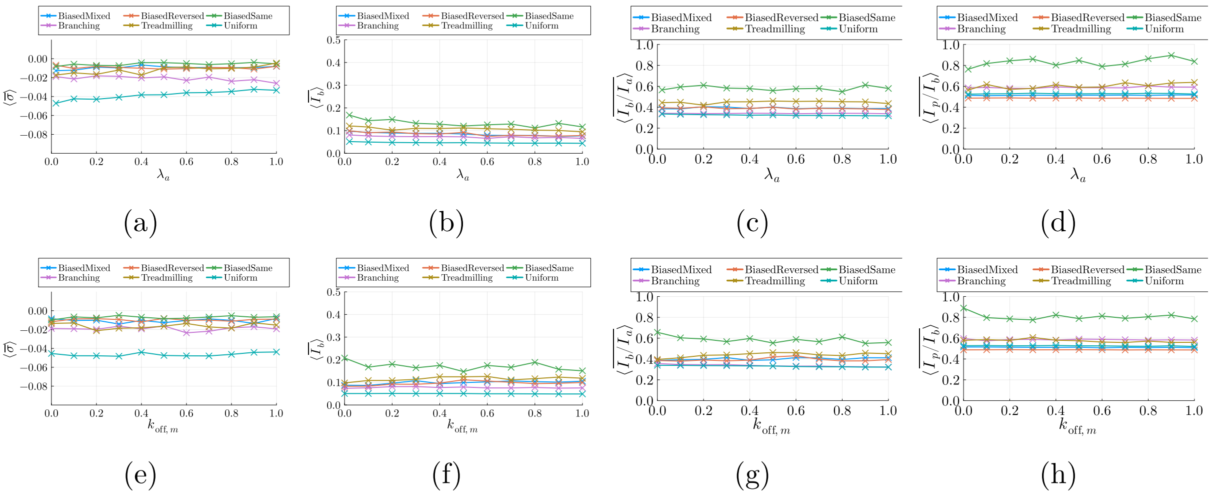


Figure B.7: The impact of varying drag and motor detachment rate on mean normal stress and bundle formation for different turnover models.
